# Memory-based and surround-based pattern completion show distinct laminar profiles in human V1

**DOI:** 10.64898/2026.09.24.753595

**Authors:** Zirui Zhang, Clement Abbatecola, Yingying Huang, Yulia Lazarova, Angus T. Paton, Maria Wimber, Christiaan N. Levelt, Matthew E. Larkum, Lars F. Muckli, Lucy S. Petro

## Abstract

Visual input is often incomplete, requiring the brain to infer missing information from prior experience or contextual cues. Establishing whether memory-derived and context-derived information shape pattern completion through shared or distinct top-down pathways is central to understanding how perception emerges from the integration of sensory input and prior knowledge. Across two experiments, we used 7T layer fMRI to measure activity in an occluded region of human V1 while participants viewed natural scenes. Memory-based completion was enabled by varying the level of exposure to full scene images before scanning, while surround-based completion was implemented via two scenes that were repeatedly presented in partially occluded form but never seen intact. In both fMRI experiments, category decoding in the occluded V1 region only emerged after multiple prior exposures, whereas novel scenes did not show reliable decoding. Repeated partially occluded scenes also supported decoding, despite the hidden region never being shown, consistent with surround-based pattern completion. Critically, mnemonic and surround-based pattern completion showed opposite laminar decoding profiles across V1 depths, with mnemonic feedback dominating deep layers, and surround-based feedback dominating superficial layers. Together, these findings suggest that completion in the early visual cortex depends on the sources of inferred information, revealing dissociable feedback pathways for memory- and context-driven inference.

## Introduction

Recognising a familiar scene is possible even when visual input is incomplete. When part of an image is occluded, the brain infers what is missing by combining current contextual information with prior knowledge, a process that is central to perception in natural environments where sensory input is typically partial, ambiguous or degraded. The ability to recover a complete representation from incomplete input, pattern completion, has traditionally been linked to functions of the medial temporal lobe, particularly the hippocampus, and to higher-order visual representations that support invariant object and scene identity^1,2,3^. However, it remains unclear how completed representations are expressed across the cortical hierarchy, and whether they can influence the earliest stages of visual processing. Resolving this question would advance our mechanistic understanding of how top-down information is integrated within sensory circuits to construct perceptual experience.

The primary visual cortex (V1) provides a critical test of whether memory-based information influences early sensory representations during recognition. V1 is retinotopically organised and strongly driven by visual input^4^. Thus, when part of a scene is physically occluded, information about the missing content is not available to the corresponding retinotopic region of V1 through feedforward sensory input. However, V1 also receives extensive feedback from higher visual and associative areas, providing a pathway through which contextual or mnemonic information could support the inference of missing content and shape early visual representations^5,6,7,8,9,10^.

Visual cortical activity has been observed in the absence of sensory input, including during imagery, prediction, and memory retrieval^11,12,13,14,15,16,17,18,19,20,21,22^. However, the distinction between mnemonic and surround-driven pattern completion in the primary visual cortex remains unclear, limiting our understanding of whether perceptual completion relies on reinstating image-specific information from previous encounters or on predictions derived from the current visual context and learned statistical regularities of the world. Although both forms of completion therefore depend on prior experience, they draw on different forms of information. Missing content may be inferred from generic visual regularities, such as local context, contour continuation, or scene structure, allowing surround completion even for novel stimuli without prior experience of the exact image. Alternatively, when an occluded image is familiar, recognition may recruit stored information about that specific image, allowing missing content to be reinstated from memory. These processes may both involve top-down signals, but they differ in their computational mechanisms: one generates plausible structure by combining the currently available visual context with learned statistical regularities, whereas the other reinstates information associated with a specific prior encounter. A key unresolved question is whether memory-based information can project back into the early visual cortex in a way that is distinct from surround-based pattern completion.

Here we used high-resolution laminar fMRI to test how prior experience shapes representations in human V1 during scene recognition under occlusion. Participants viewed scene images in which a critical region was occluded, while prior exposure to the full image was manipulated, allowing us to separate information available from current sensory input from information available through memory. We asked whether category-specific information could be decoded from the occluded retinotopic region of V1, whether this information depended on familiarity, and whether memory-guided reinstatement could be distinguished from surround-based completion across cortical depths. We found that V1 contributes to recognition in a task-dependent and layer-specific manner. When recognition required inference about occluded content, highly familiar images elicited category-specific information in deep layers. In contrast, surround pattern completion was associated with a distinct superficial-layer profile and did not depend on prior exposure.

## Results

We used ultra-high-field 7 Tesla functional magnetic resonance imaging (fMRI) to investigate how top-down contextual signals in the primary visual cortex are modified by memory in 41 participants (32 after data quality control) across two experiments (Fig. 1a).

**Fig. 1.**
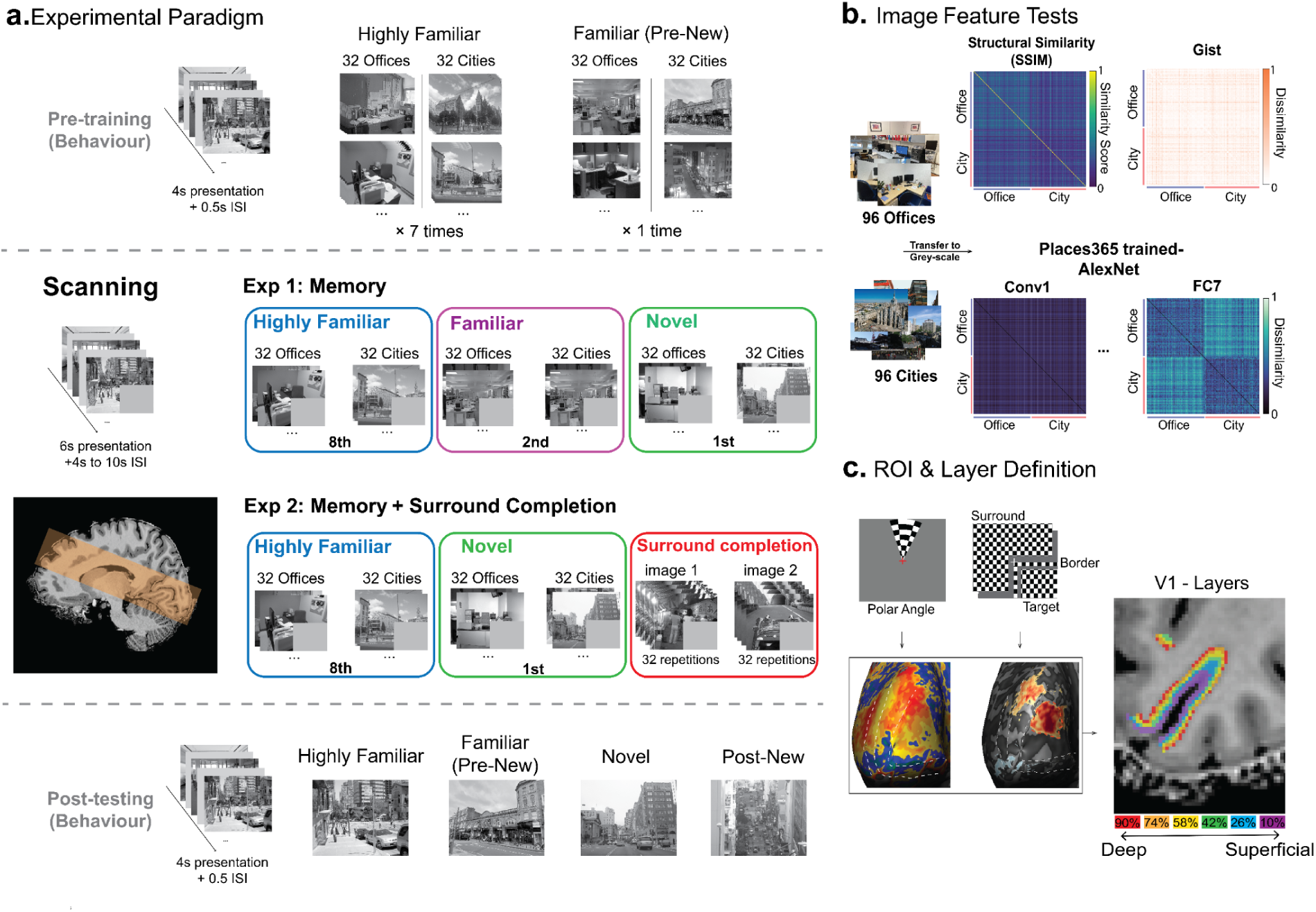
Experimental design and laminar ROI definition in V1. **a.** Experimental procedure. Each experiment consisted of pre-training (non-occluded images), fMRI scanning and post-scan behavioural testing. During scanning in Experiment 1, participants (N = 18) viewed natural scene images with a lower-right occluder under three familiarity conditions: Novel images (green), which had not been seen before scanning; Familiar images (purple), which had been seen once during pre-training; and Highly Familiar images (blue), which had been seen seven times during pre-training. Each condition contained 64 images, comprising 32 city and 32 office scenes, and we presented each image once during scanning. Experiment 2 (N = 14) followed the same general procedure but included only the Novel and Highly Familiar conditions, with an additional surround-completion condition. In this condition, we repeatedly presented two partially occluded images 32 times each (never shown in their full, unoccluded form), matching the number of presentations in each familiarity-by-category condition. After scanning, participants completed a recognition-memory test (post-test). We randomly selected sixteen images from each familiarity condition, including eight city and eight office scenes, and presented them together with 16 new images that served as foils. Participants performed an OLD/NEW recognition task, allowing memory performance to be quantified separately for each condition. **b.** Image feature analysis. The stimulus set comprised 192 natural scene images, including 96 city and 96 office scenes, all converted to grayscale. We performed image feature analyses to reduce potential low-level confounds. We quantified structural similarity between images using the Structural Similarity Index Measure (SSIM), and highly similar images were replaced to minimise image-specific similarity effects on memory performance. We constructed representational dissimilarity matrices (RDMs) from GIST descriptors and AlexNet features using 1 − Pearson’s r as the dissimilarity metric. GIST RDMs were used to quantify global scene-layout similarity across images, whereas AlexNet RDMs, derived from the early convolutional to late fully connected layers of a Places365-trained network, were used to verify that city and office scenes remained separable in the model’s feature space. **c.** Retinotopic ROI and cortical-depth definition. We used retinotopic checkerboard mapping to define the occluded (target) region in V1 and to separate it from what we term the border and surround regions. The region we refer to as the target region corresponded to the lower-right quadrant of the visual field, the border region captured the transition zone around the occluder, and the surround region corresponded to the remaining stimulated visual field. We used a GLM contrast comparing target activity with border and surround activity to delineate the V1 ROI receiving feedback signals. We used independent polar-angle mapping with high-contrast flickering checkerboards to define the boundaries of early visual areas V1–V3. The left occipital cortical reconstruction shows the target region (occluded area) within V1, outlined in red. We defined cortical depth by segmenting the grey matter into six equivolume depth levels, corresponding to 10%, 26%, 42%, 58%, 74% and 90% cortical depth, from superficial grey matter (near the cerebrospinal fluid) to deep grey matter (near the white matter). We used these depth levels to examine laminar profiles of decoding and activity across V1.

In Experiment 1 (N = 18), we measured memory-dependent contextual feedback signals. Participants viewed full-scene images from two categories (city and office) that were each shown exactly once in the 7T scanner, but differed in prior exposure levels: Highly Familiar (pre-trained seven times before brain scanning), Familiar (pre-trained once), and Novel (untrained, seen for the first time during brain scanning). During the fMRI session, we occluded the lower-right quadrant of all 192 images, masking feedforward inputs to this retinotopic region of V1, and therefore enabling us to isolate feedback-driven contextual signals. For comparison, we defined a stimulated V1 control region in the right hemisphere, corresponding to the contralateral, non-occluded portion of the image and therefore driven by direct feedforward visual input. This feedforward region also receives feedback signals, but for simplicity, we refer to it as our feedforward region.

In Experiment 2 (N = 14), we compared these memory-driven feedback signals to those involved in surround completion, meaning a cortical representation of the image continuing in space over the occluded region, even in the absence of feedforward information in that region or related memory signal^15^. To isolate top-down contextual inference from memory reinstatement, participants were repeatedly presented with two unique, partially occluded scenes (a market and a car scene; ^11^) that were never shown in full. This ensured that the missing information was inferred dynamically from the visible surround rather than reconstructed by retrieving a previously encoded full-image template. We intermixed these two stimuli with Highly Familiar and Novel unrepeated categorical images (identical to the ‘Novel’ and ‘Highly Familiar’ conditions in Experiment 1), allowing direct comparison of memory-based versus surround completion signals in layers of V1 within the same scanning session (Fig. 1a).

In both experiments, participants performed an OLD/NEW recognition task to assess memory performance during scanning. We performed retinotopic mapping procedures to define the lower-right target region and delineate the boundaries of V1, V2, and V3 (Fig. 1c). We used multivoxel pattern analysis (MVPA) to quantify category decoding accuracy across memory levels and to evaluate feedback-related decoding associated with surround completion. We applied a linear mixed-effects model to assess trends across cortical depths associated with distinct types of feedback signals.

### Memory-Enhanced Categorical Feedback Decoding Shows Opposite Layer Profiles to Surround Completion Decoding in V1

We first explored the effect of memory on contextual feedback processing in V1. To this end, we performed category decoding in layers of the non-stimulated V1 area for each memory condition. No item-level classifiers could be used here since we presented each image only once in the scanner. We trained participants prior to scanning by presenting images in nonoccluded form 7 times (Highly Familiar) or once (Familiar) to build varying levels of familiarity with the full scene, while images in the Novel condition were untrained. During the fMRI sessions, participants judged each image as ‘old’ or ‘new’ to engage memory.

Categorical decoding (City vs Office) was absent in the occluded region of interest for Novel and Familiar images, with no cortical depth showing significant above-chance performance (all FDR-corrected *ps* ≥ 0.181) (Fig. 2a). In contrast, Highly Familiar images were significantly decoded at 90% Cortical depth (deepest layer, Accuracy = 0.58, CI [0.550, 0.607], FDR-corrected p < 0.001), 74% ( second deepest layer, Accuracy = 0.56, CI [0.529, 0.588], FDR-corrected p < 0.001), 58% (middle layer, Accuracy = 0.55, CI [0.526, 0.578], FDR-corrected p < 0.001), and 42% (middle layer, Accuracy = 0.54, CI [0.515, 0.570], FDR-corrected p < 0.01), whereas decoding at two superficial layers (cortical depth 26% and 10%) did not differ significantly from chance level (Fig. 2a). These findings indicate that category information in the feedback region emerged only after extensive visual experience, suggesting that memory-related feedback enhanced the representation of categorical information in the occluded region of V1.

**Fig. 2.**
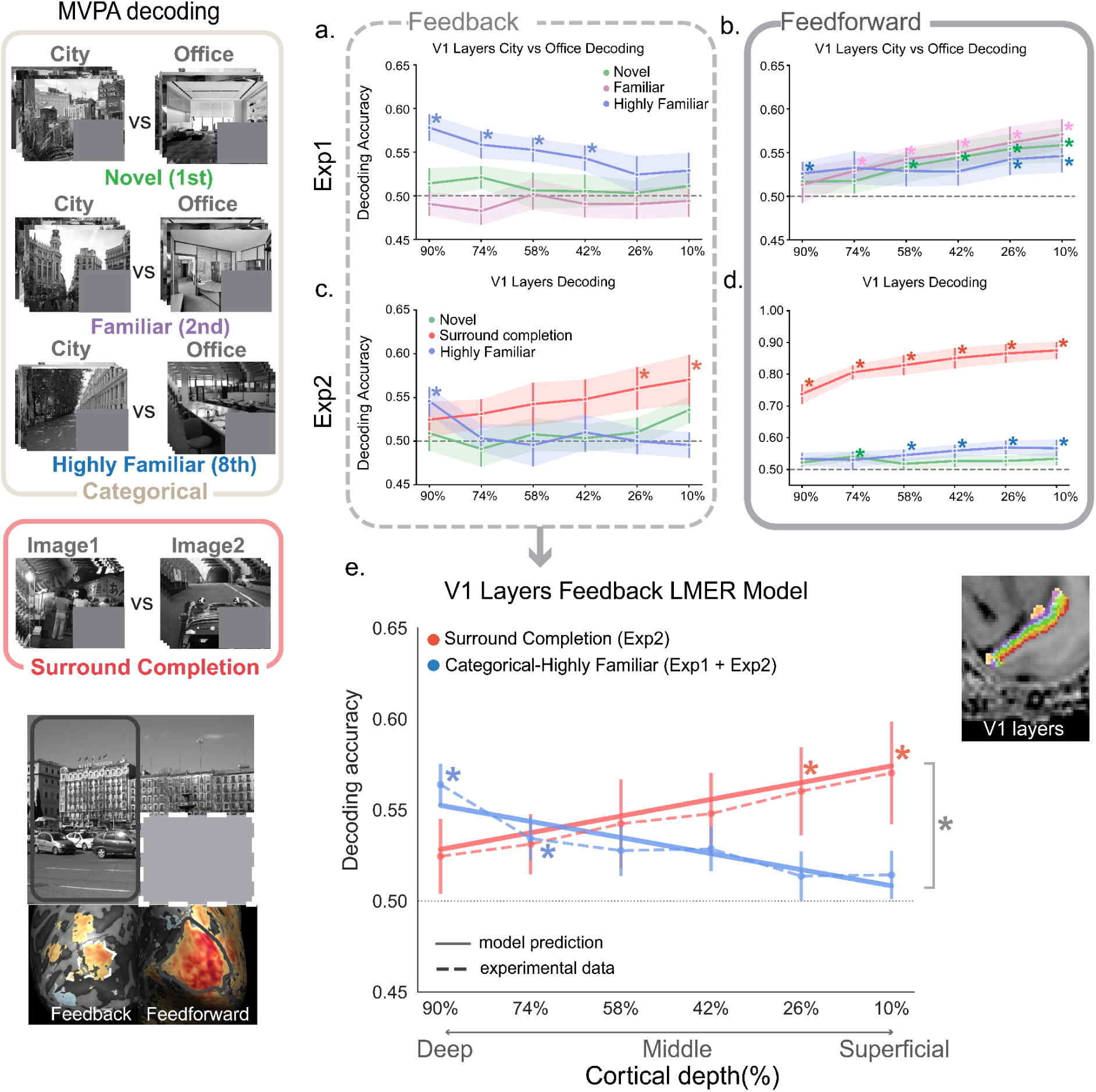
Laminar profiles of decoding in V1 depend on memory and surround completion conditions. **a,b.** City–office decoding accuracy across cortical depth in Experiment 1 (N = 18) for the V1 feedback region (**a**) and V1 feedforward region (**b**), shown for Novel, Familiar and Highly Familiar scenes. **c,d.** City–office decoding accuracy in Experiment 2 (N = 14) for Novel and Highly Familiar scenes, and image-specific decoding for the surround-completion condition, shown for the V1 feedback region (**c**) and V1 feedforward region (**d**). **e.** Linear mixed-effects model comparing the laminar decoding profiles of Highly Familiar scenes across Experiments 1 and 2 (N = 32) with surround-completion decoding from Experiment 2 (N = 14). Dashed lines show observed decoding accuracy and solid lines show model-predicted values. Cortical depth is ordered from deep to superficial layers. The horizontal line indicates the chance level at 0.50. Asterisks show statistically significant decoding above chance or significant model effects, as indicated in the figure. Error bars denote the standard error of the mean (SEM). *p < 0.05, Benjamini–Hochberg false discovery rate corrected.

In Experiment 1, category decoding required the classifier to identify information shared across multiple examples of the same scene category. Because each example had a different visible surround region, the surround did not uniquely predict the specific missing content. Decoding in the occluded V1 region could therefore reflect a combination of category-consistent information in the visible surround and memory-based reinstatement of previously viewed scene content. By contrast, in the repeated-image condition, the same visible surround was repeatedly paired with the same occluded region. This stable image-specific association made the surround a highly reliable cue to the missing content, allowing the representation in occluded V1 to be supported by surround-based pattern completion. To dissociate these two mechanisms, we conducted Experiment 2 by incorporating a repeated-image condition similar to that used by Muckli et al.^15^, allowing us to directly compare memory-dependent category feedback with surround-driven image feedback. Consistent with Experiment 1, categorical decoding of Novel images remained absent across all cortical depths in the V1 feedback region. In contrast, Highly Familiar images showed significant category decoding at 90% cortical depth (deepest layer; Accuracy = 0.55, CI [0.515, 0.577], FDR-corrected p < 0.05) (Fig. 2c), replicating the deep layer decoding profile that we observed in Experiment 1. Importantly, under surround completion conditions, participants never saw the lower-right occluded quadrant, meaning that feedback information could only be inferred from the repeatedly presented surrounding image context. Under these conditions, decoding of the repeated single images was significant at 26% cortical depth (second superficial layer, Accuracy = 0.56, CI [0.514, 0.605], FDR-corrected p < 0.05) and 10% cortical depth (superficial layer, Accuracy = 0.57, CI [0.518, 0.624], FDR-corrected p < 0.05) (Fig. 2c), replicating the superficial layer profile that we previously reported^15^.

To test consistency across experiments, we fitted a linear mixed-effects model predicting decoding accuracy from condition, cortical layer as a linear variable, experiment, ROI, and their interactions, with participants included as a random intercept. We restricted the analysis to the three conditions common to both experiments (categorical-Novel, categorical-Highly Familiar and Novel-vs-Highly Familiar) There was no significant main effect of experiment, [F(1, 47.46) = 0.48, p = 0.49], and no significant interactions between experiment and condition, [F(2, 1098) = 1.16, p = 0.32], experiment and layer, [F(1, 1098) = 0.07, p = 0.80], or condition, layer, and experiment, [F(2, 1098) = 0.71, p = 0.49]. Laminar-trend analyses also showed no significant experiment differences within each condition and ROI (all ps> 0.05; Supplementary Fig. 6), indicating that laminar decoding profiles were consistent across experiments. These results demonstrate that the laminar decoding profiles were highly consistent across independent experiments, indicating that the observed layer-specific effects reflect robust and reproducible cortical processing mechanisms rather than experiment-specific influences. These results allowed us to proceed with the analysis of decoding profiles by condition while accumulating evidence across experiments.

Pooled categorical decoding analyses revealed significant Highly Familiar category decoding in the V1 feedback region at 90% cortical depth (deepest layer, Accuracy = 0.56, CI [0.542, 0.586], FDR-corrected p < 0.001) and 74% cortical depth (second deepest layer, Accuracy = 0.53, CI [0.511, 0.557], FDR-corrected p < 0.05; Fig. 2e), whereas Novel images did not show significant decoding at any cortical depth (all FDR-corrected ps > 0.13). We further compared decoding accuracy between Highly Familiar categorical decoding and surround completion decoding using a linear mixed-effects model with condition, cortical Layer, ROI, and their interactions as fixed effects, and participant included as a random intercept. This analysis revealed significant main effects of Condition, [F(1, 531.05) = 78.77, p < 0.001], and Layer, [F(1, 512.38) = 16.57, p < 0.001], as well as a significant Condition × Layer interaction, [F(1, 512.38) = 23.60, p < 0.001]. Follow-up laminar trend analyses showed that Highly Familiar categorical decoding significantly decreased from deep to upper cortical depths [t(513) = -2.93, p < 0.01], whereas surround completion decoding significantly increased towards the upper cortical depth [t(513) = 2.02, p < 0.05]. Direct comparison of these trends showed a significant difference between the two conditions [t(513) = 3.30, p = 0.001] (Fig. 2e). Overall, we find that memory-based and surround-based feedback signals are expressed through distinct laminar profiles in V1.

### Behavioural testing showed graded recognition memory across conditions

Before assessing how prior experience shaped laminar V1 responses, we first verified that participants recognised the trained images while correctly rejecting novel images. Participants were trained to remember images from the Highly Familiar condition during the pre-training session, and we assessed memory performance during scanning using the same recognition task. The behavioural results showed that participants demonstrated clear memory for images presented seven times during training, and images in the ‘Novel’ condition were recognised as ‘have seen before’ at a lower rate (Supplementary Fig. 10 and Note 1).

Images in the Familiar condition, which had been presented once during pre-training and were encountered for the second time during scanning, were recognised less often than Highly Familiar images. For single-repeated images in the Surround completion condition, “seen before” responses increased with repetition and ultimately exceeded those for Highly Familiar images (Fig. 3a). Consistent with this pattern, scanning-session behavioural responses differed significantly across familiarity conditions (Kruskal–Wallis test, H(3) = 216.22, p < 0.001). Post-hoc Dunn’s tests with Benjamini–Hochberg correction revealed a graded increase in “seen before” responses across conditions: Novel < Familiar < Highly Familiar < Surround completion. Participants responded ‘seen before’ significantly less often for Novel images than for either repeated-image condition (all ps < 0.001, FDR-corrected); Familiar images were endorsed less often than both Highly Familiar and Surround completion images (both ps < 0.001, FDR-corrected), and Highly Familiar images were endorsed less often than Surround completion images (p < 0.05, FDR-corrected). Reaction times showed a different pattern from recognition-response rates (Fig. 3b), but also varied significantly across conditions, H(3) = 420.94, p < 0.001. Post-hoc Dunn’s tests showed significant reaction-time differences between most condition pairs, including Novel versus Highly Familiar, Novel versus Surround completion, Familiar versus Highly Familiar, Familiar versus Surround completion, and Highly Familiar versus Surround completion (all ps < 0.001, FDR-corrected), whereas Novel and Familiar images did not differ significantly, p = 0.053. Finally, post-testing further confirmed that participants formed distinct levels of familiarity across conditions in both experiments (Supplementary Fig. 10 and Note 1).

**Fig. 3.**
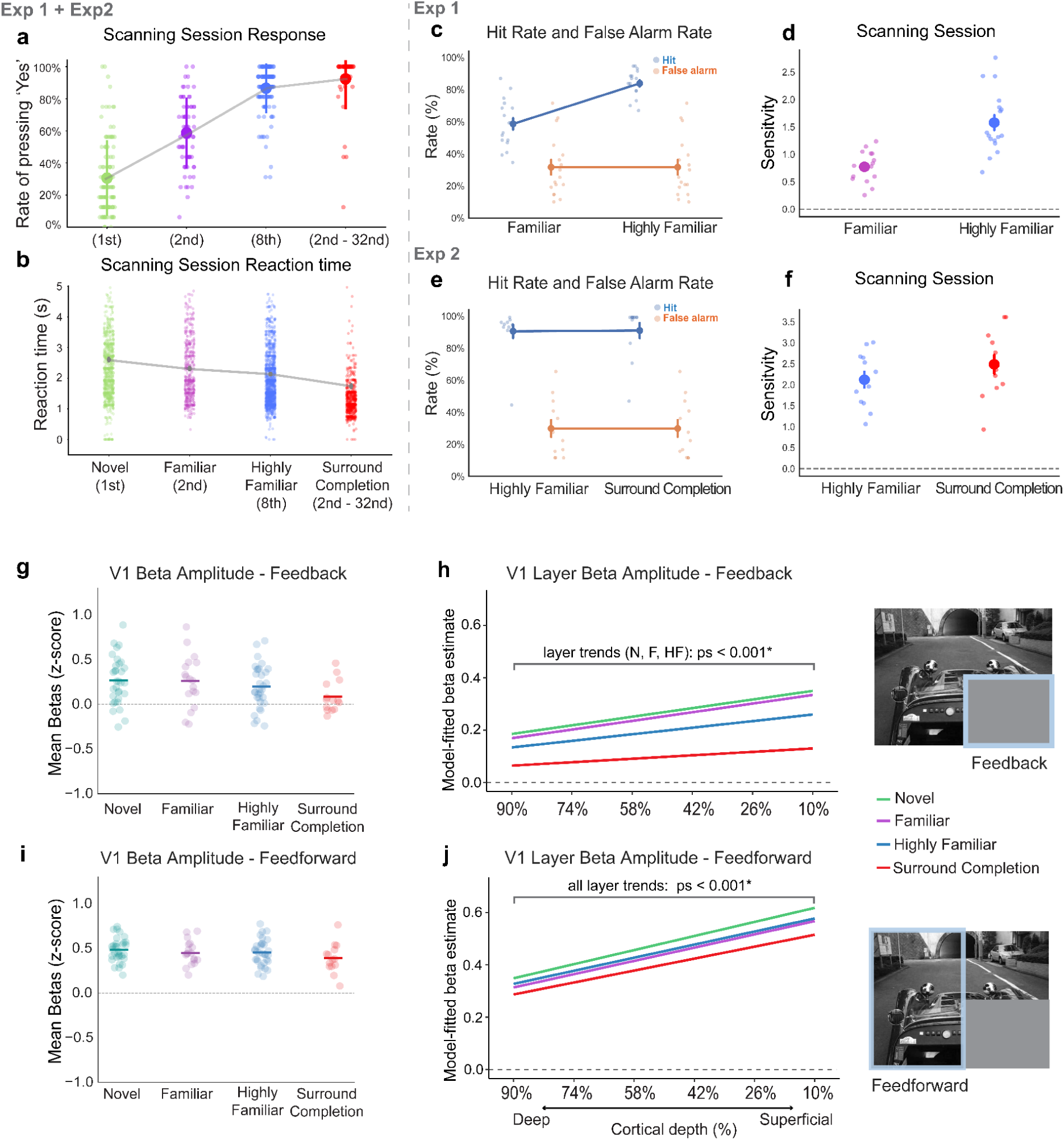
Behavioural performance and univariate BOLD responses. **a,b.** Behavioural performance during the scanning session. **a.** Proportion of “yes, I have seen this image” responses across conditions, including Novel images (green; Experiments 1 and 2, N = 29, excluded 3 participants from both experiments due to technical problems), Highly Familiar images (blue; Experiments 1 and 2), Familiar images (purple; Experiment 1 only, N =17), and Surround completion trials from the 2nd to 32nd presentation (Experiment 2 only, N = 12). For Novel images, “yes” responses correspond to false alarms, whereas for Familiar, Highly Familiar, and Surround completion trials they correspond to hits. Error bars denote the standard deviation (SD). **b.** Reaction times for the same conditions during the scanning session. Error bars denote the standard error of the mean (SEM). **c,d.** Recognition-memory performance in Experiment 1 (N = 17). Error bars denote the standard error of the mean (SEM). **c.** False-alarm rate for Novel images and hit rates for Familiar and Highly Familiar images. **d.** Signal-detection sensitivity (SDT), quantified as d′, for Experiment 1, where Novel images served as the noise condition and Familiar/Highly Familiar images served as signal conditions. **e,f.** Recognition-memory performance in Experiment 2 (N = 12). Error bars denote the standard error of the mean (SEM). **e.** False-alarm rate for Novel images and hit rates for Highly Familiar and Surround completion trials. **f.** Signal-detection sensitivity, quantified as d′, for Experiment 2, where Novel images served as the noise condition and Highly Familiar/Surround completion trials served as signal conditions. **g, i.** Mean BOLD signal changes (z-transformed) across conditions in the **g,** V1 feedback ROI and **i,** the V1 feedforward ROI. Conditions include Novel and Highly Familiar trials from Experiments 1 and 2 (N = 32), Familiar trials from Experiment 1 (N = 18), and Surround completion trials from Experiment 2 (N = 14). Dots indicate individual participants, and horizontal lines indicate the condition means. **h, j.** Model-fitted beta estimates across cortical depth for Novel, Highly Familiar, and Surround completion conditions, shown separately for **h,** the V1 feedback ROI and **j,** the V1 feedforward ROI. Lines indicate fixed-effect predictions from the linear mixed-effects model. All *p < 0.05, Benjamini–Hochberg false discovery rate corrected.

Signal-detection analysis revealed robust recognition memory across both experiments, with participants reliably discriminating previously seen images from novel foils. Recognition sensitivity was quantified by treating Novel images as the noise distribution: “yes” responses to Novel images were counted as false alarms, whereas “yes” responses to Familiar, Highly Familiar and Surround completion images (from the second presentation) were counted as hits. In Experiment 1, recognition performance increased with prior exposure. Highly Familiar images were recognised more reliably than Familiar images, with higher hit rates (median: Familiar = 0.59; Highly Familiar = 0.84; paired t(16) = 7.54, p < 0.001) and higher recognition sensitivity (Wilcoxon signed-rank test, W = 0.00, p < 0.001). Median sensitivity increased from 0.78 for Familiar images to 1.34 for Highly Familiar images (Fig. 3c,d). These results indicate that repeated pre-training strengthened recognition memory. In Experiment 2, Surround completion images showed recognition sensitivity comparable to Highly Familiar images [paired t(11) = 1.18, p = 0.262], and hit rates did not differ significantly between the two conditions (median: Surround completion = 0.99; Highly Familiar = 0.95; Wilcoxon signed-rank test, W = 31.00, p = 0.569). Thus, repeated exposure in the Surround completion condition produced recognition performance comparable to that observed for Highly Familiar images.

### Repeated exposure suppresses univariate BOLD responses in V1

To test whether familiarity altered BOLD response amplitude, we fitted a linear mixed-effects model with condition, cortical layer and ROI as fixed effects, and participant as a random intercept. This analysis revealed significant main effects of condition, F(3, 1107.5) = 5.92, p < 0.001, layer, F(1, 1105.0) = 249.12, p < 0.001, and ROI, F(1, 1105.0) = 154.22, p < 0.001. We also found a significant layer × ROI interaction, F(1, 1105.0) = 24.86, p < 0.001, indicating that laminar response profiles differed between feedback and feedforward V1. In contrast, the condition × layer, condition × ROI and condition × layer × ROI interactions were not significant (all ps ≥ 0.237), suggesting that condition effects were broadly similar across cortical depths and ROIs. Overall response amplitudes differed reliably between conditions, with higher amplitudes in the feedforward ROI than in the feedback area. In the feedback ROI, response amplitudes were highest for Novel and Familiar images, with no significant difference between these conditions (p = 0.320). Both showed greater responses than Highly Familiar and Surround Completion trials, while Highly Familiar responses also exceeded Surround Completion responses (all FDR-corrected ps ≤ 0.001; Fig. 3g). In the feedforward area, responses on Novel images were higher than Highly Familiar, Familiar and Surround Completion responses (all FDR-corrected ps ≤ 0.034), and Highly Familiar responses exceeded Surround Completion responses (p = 0.013); the remaining comparisons were not significant (Fig. 3i). Response amplitudes increased from deep to superficial cortical layers for Novel, Familiar and Highly Familiar conditions in both ROIs (all ps < 0.001), and for Surround Completion in the feedforward ROI (p < 0.001), whereas the laminar gradient for Surround Completion was not significant in the feedback ROI (p = 0.115; Fig. 3h,j). Importantly, however, pairwise comparisons revealed no significant differences in laminar gradients between conditions within either ROI after FDR correction (feedback, all ps ≥ 0.228; feedforward, all ps ≥ 0.878). These results indicate that repeated exposure reduced overall V1 response amplitudes, while the shape of the laminar response profile remained largely stable across familiarity conditions.

To further test whether repeated presentation caused adaptation in the surround-completion condition, we fitted another linear mixed-effects model including Run (1-4), Cortical depth (six cortical depths), ROI (V1 feedback/feedforward), and their interactions, with participants as a random intercept. There was no main effect of Run and no Run-related interactions (all ps > 0.26). In the feedback ROI, activity did not differ across runs (all FDR-corrected ps ≥ 0.159, Supplementary Fig. 9a), whereas in the feedforward ROI, Run 1 (first run) showed higher activity than Runs 2–4 (all FDR-corrected ps ≤ 0.0004), with no differences among Runs 2–4 (all ps > 0.26, Supplementary Fig. 9b). Laminar profiles also remained stable across runs. The feedback ROI showed no significant laminar trend in any run (all ps = 0.195, Supplementary Fig. 9c), whereas the feedforward ROI showed a significant increase in activity from deep to superficial layers in all four runs (all ps < 0.0001, Supplementary Fig. 9d). Importantly, laminar slopes did not differ across runs in either the feedback (all ps = 0.966) or feedforward ROI (all ps > 0.76). The low response amplitude in the surround-completion condition was not explained by progressive adaptation across runs in the feedback ROI, while the feedforward ROI showed only an initial reduction followed by stable response amplitudes and laminar profiles.

### Memory-related feedback alters low-level feature representations in V1

In complex natural scene stimuli, multiple features coexist within an image and are processed at different stages of the visual hierarchy. Although our decoding analyses showed that prior experience enhances category information in occluded V1, they do not indicate whether this familiarity-related representation reflects low-level image structure, higher-level scene information, or a combination of both.

The primary visual cortex is primarily responsible for encoding low-level visual features at the earliest stage of visual processing^23,24,25,26,27^. Convolutional neural networks (CNNs), particularly standard feedforward architectures trained for visual recognition, capture key aspects of hierarchical visual processing and provide a valuable framework for testing representational correspondences between human fMRI responses and features from different network layers^28,29,30,31^. AlexNet provides a relatively simple hierarchical model in which earlier convolutional layers capture lower-level visual structure and later fully connected layers capture more abstract scene-related representations^32,33,34^. We constructed 8 × 8 block-averaged representational dissimilarity matrices (RDMs) across categories and experimental runs for each familiarity condition, and then assessed Spearman’s rank-order correlation between fMRI representations within V1 layers and AlexNet layers (Fig. 4a).

**Fig 4.**
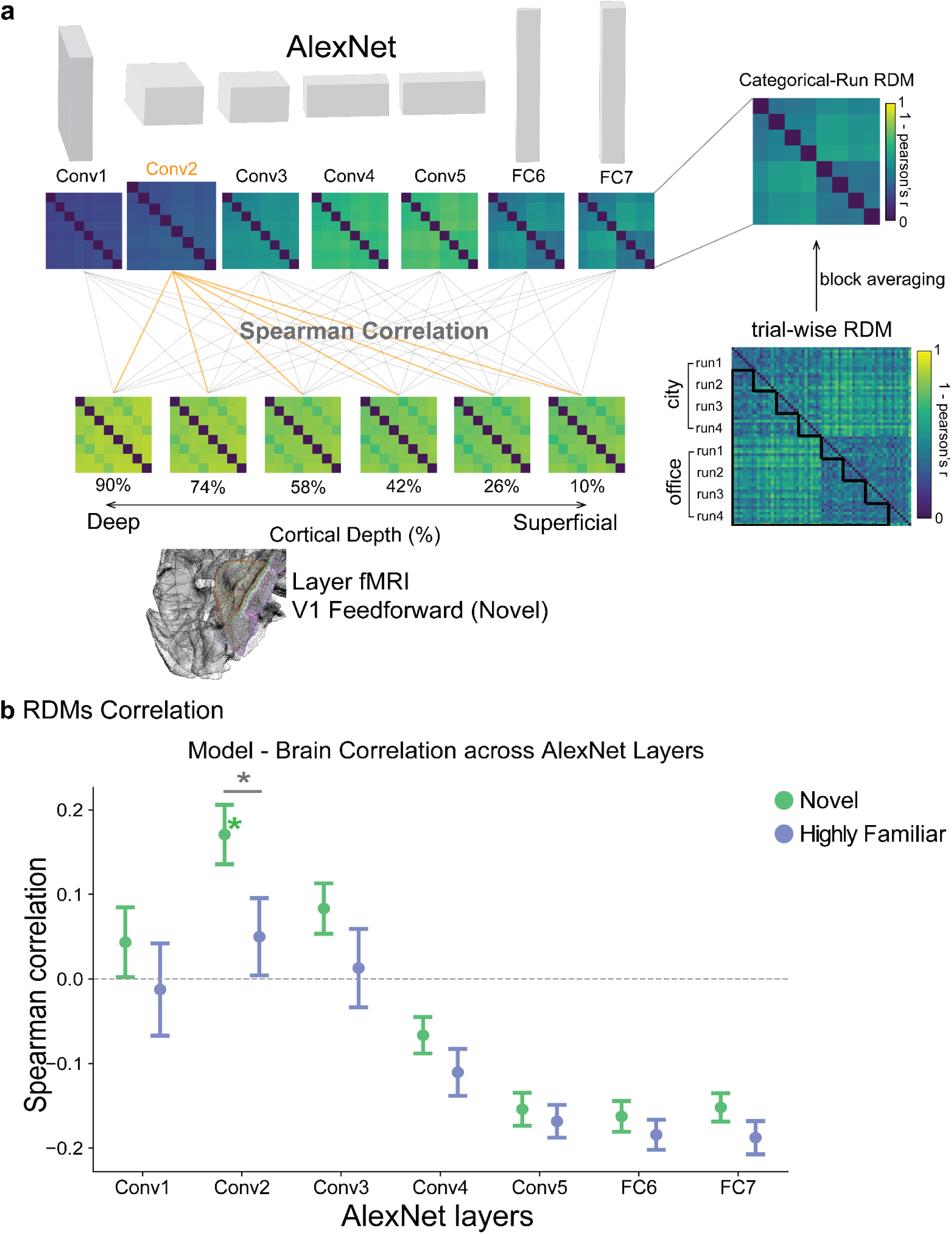
fMRI and AlexNet RDMs correlation. **a.** Example plots illustrate the block-averaging procedure from trial-wise representational dissimilarity matrices (1-Pearson’s r RDMs) to categorical-run RDMs for both fMRI and AlexNet layer RDMs for one participant. Spearman rank correlations were computed for each pair of fMRI and AlexNet-layer RDMs within each familiarity condition. The example plot shows the RDMs correlation in the Novel condition between the V1 feedforward area and AlexNet layers. **b.** RDMs correlation results showing representational similarity between fMRI and the AlexNet model across familiarity conditions for both experiments (N = 32). The analysis compares Novel and Highly Familiar conditions using Spearman rank correlations from the Conv1 layer to the FC7 layer with all V1 layers. Conv2 features showed significant correlations with all layers of V1 feedforward data (see Supplementary Fig. 5b), whereas the Highly Familiar condition showed no significant correlation. We assessed statistical significance using one-sided t-tests against a null correlation of zero (p < 0.05, FDR-corrected). In contrast, other higher-level layers showed no significant correlation or difference between the Highly Familiar and Novel conditions. Linear mixed-effects model comparing fMRI–AlexNet correlations between Novel and Highly Familiar conditions across two experiments (N = 32). Error bars denote the standard error of the mean (SEM). *p < 0.05, Benjamini–Hochberg false discovery rate corrected.

In the feedforward area, representational similarity in the Novel condition was significantly correlated with AlexNet Conv2 across all cortical depths (Spearman’s R = 0.163–0.185; all FDR-corrected ps < 0.008, one-sample right-tailed t-tests against zero; Supplementary Fig. 5b). No other condition showed significant correlations with any AlexNet convolutional layer in either the feedforward or feedback ROI (all FDR-corrected ps > 0.05). For further comparisons, we fitted a model of the correlation r-value with two conditions (Novel and Highly Familiar) × six cortical depths (fMRI layers) × seven AlexNet layers × two ROIs (V1 Feedback and Feedforward areas), with participants as a random effect. The analysis revealed significant main effects of condition, F(1, 5289) = 7.45, p = 0.006, AlexNet layer, F(6, 5289) = 59.50, p < 0.001, and ROI, F(1, 5289) = 9.21, p = 0.002. Importantly, there was a significant condition × AlexNet layer interaction, F(6, 5289) = 4.56, p < 0.001, indicating that the pattern of fMRI correspondence across AlexNet layers differed between familiarity conditions. However, there was no significant effect on cortical depths [F(1, 5289) = 1.33, p = 0.25], nor was there a significant Condition × Layer × ROIs interaction [F(1, 5289) = 0.63, p = 0.43], suggesting that the observed effects were consistent across cortical depth and ROI. To test whether the familiarity effect was specific to lower-level rather than higher-level visual representations, we contrasted Conv2, an early feature-processing layer, with FC7, a late layer encoding more abstract visual information. Correlation values between the V1 feedforward area and AlexNet layer Conv2 under the Novel condition were significantly higher than the Highly Familiar condition in the feedforward area (p < 0.001, FDR-corrected), while the last connection layer showed no significant difference between familiarity conditions (p > 0.10, FDR-corrected) (Fig. 4b and Supplementary Fig. 5b). These findings indicate that novel visual input in V1 is more strongly associated with lower-level CNN feature representations, whereas prior experience reduces the correspondence between V1 activity and low-level image features.

### Decoding in the feedforward region of V1

Feedforward categorical decoding showed a consistent familiarity-dependent pattern across both experiments. In Experiment 1, Novel images were significantly decoded in middle and superficial layers (Accuracy: 0.534– 0.558, FDR-corrected ps < 0.05), Familiar images across all layers except the deepest layer (Accuracy: 0.529 – 0.57, FDR-corrected ps < 0.05), and Highly Familiar images in the deepest (90%) and two superficial layers (26% and 10%) (Accuracy: 0.53–0.55, FDR-corrected ps < 0.05) (Fig. 2b). In Experiment 2, Novel decoding was significant only at 74% cortical depth (second deepest layer, Accuracy = 0.54, CI [0.515, 0.571], FDR-corrected p < 0.001), whereas Highly Familiar images showed significant decoding in two middle depths (58% and 42%) and superficial depths (26% - 10%) (Accuracy: 0.54–0.57, FDR-corrected ps < 0.05). Decoding of repeatedly presented single images (surround completion) was robust across all cortical depths (0.74–0.88, all FDR-corrected ps < 0.001) (Fig. 2d). Consistent with the absence of significant experiment effects in the mixed-effects model, pooled analyses revealed significant categorical decoding across all cortical depths for both Novel (0.52–0.55) and Highly Familiar images (0.53–0.56) (all FDR-corrected ps < 0.05).

Memory decoding (Novel vs Highly Familiar) in the feedforward area was substantially weaker, reaching significance only at superficial layer cortical depth in Experiment 1 (Accuracy = 0.53, CI [0.510, 0.557], FDR-corrected p = 0.009) and Experiment 2 (Accuracy = 0.54, CI [0.512, 0.570], FDR-corrected p = 0.007), and at middle-superficial to superficial depths (42%, 26%, and 10%) in the pooled analysis (Accuracy: 0.52–0.54, FDR-corrected ps < 0.05). In the feedback region, however, we observed no significant memory decoding in Experiment 1 (Accuracy: 0.51–0.52, all FDR-corrected ps > 0.17), Experiment 2 (Accuracy: 0.50–0.52, all FDR-corrected ps > 0.27), or the pooled analysis (0.50–0.51, all FDR-corrected ps > 0.16) (Supplementary Fig. 6). These results show the V1 feedforward region showed reliable category decoding across conditions, consistent with stronger contributions from visible sensory input. Thus, memory appeared to enhance feedback-based category representations, even though direct decoding of familiarity was weak.

## Discussion

We investigated how prior experience influences feedback signals reaching V1 during recognition under occlusion. We found that category-relevant information emerged in the deep layers of occluded V1 only after extensive visual experience, whereas highly informative surround information supported image-specific decoding in the superficial layers of V1 even without prior exposure to the occluded content. These findings indicate that V1 receives at least two forms of top-down information: memory-derived predictions that support category-relevant inference and surround-derived predictions that support image pattern completion. When surround information alone is insufficient to specify missing content, recognition depends more strongly on memory-related feedback; when contextual information is highly predictive, feedback can be generated directly from the visible image structure.

The laminar dissociation suggests that the two types of predictions may rely on distinct feedback pathways: deep-layer feedback for memory-derived information, superficial-layer feedback for contextual completion based on currently available information. Although laminar fMRI cannot identify the precise anatomical origins of these signals, the observed dissociation is consistent with anatomical and physiological evidence showing that feedback projections terminate preferentially within superficial and deep cortical layers and may convey distinct forms of information to early sensory cortex^35,36,37^.

A central finding is that memory shaped representations in the portion of V1 corresponding to unseen image content: category-relevant information emerged in occluded V1 only after extensive prior experience, indicating that early visual cortex can receive memory-derived predictions about missing scene content. This finding is notable because early visual cortical representations are often dominated by image-specific visual structure and may show limited generalisation across examples from the same category^38^. Consistent with this view, category decoding was weak in the Novel condition within the feedback region, despite the presence of visible surround information. The emergence of category-relevant information after extensive experience therefore suggests that repeated exposure changed the information expressed in occluded V1, rather than simply revealing category structure already available from the visible image. Critically, after extensive visual experience, category information emerged reliably within this same retinotopic region of V1, suggesting that the effect was not driven by local feedforward input or generic surround-based completion alone. Instead, it is consistent with a model in which higher-level memory representations reinstate structured or abstract predictions about missing scene content, projecting category-relevant information back to early visual cortex in a retinotopically organised format^16,39,40^. This reinstatement may be coordinated by the hippocampus and adjacent medial temporal lobe (MTL) regions, though unlikely via direct hippocampal-V1 feedback. Computational models^41,42^, and a wealth of empirical findings^43,44^, suggest that the hippocampus reinstates distributed neocortical patterns corresponding to past experience, sometimes reaching down to V1^14,45^. Rather than projecting directly to V1, hippocampally recovered information may therefore be conveyed through a hierarchy of MTL and higher-order visual regions^46^, becoming translated into retinotopically organised predictions as it reaches the early visual cortex. Although our data cannot establish the origin or direction of this feedback, an MTL-mediated reinstatement cascade provides a plausible mechanism for the experience-dependent category information expressed in deep V1.

The laminar profile further supports this interpretation: memory-related category information was strongest in deep layers of the feedback region, whereas repeated single-image decoding was strongest in superficial layers, suggesting distinct feedback routes for memory-derived reinstatement and surround-based amodal completion. Consistent with our data suggesting that deep layers receive category-specific information derived from memory, in contrast to superficial layers which process surround completion information, investigations using neuronal recordings identify distinct functional roles for neurons in superficial and deep cortical layers (Fig. 5). For example, layer 5 pyramidal neurons (L5) do not compute sensory mismatch in the same manner as layer 2/3 neurons^47^, while L5 neurons appear to integrate information from L2/3 neurons when mice are presented with both occluded and fully visible images. Together, these findings suggest a division of labour across cortical layers in the integration of bottom-up sensory and top-down contextual information, with the evaluation of sensory mismatch in superficial layers and the processing of task relevance in deep layers. Further functional specialisation has been identified among the two major L5 pyramidal cell populations: L5 intratelencephalic (IT) neurons are more strongly informed by sensory input, whereas L5 extratelencephalic (ET) neurons receive greater information about task relevance^48^. Importantly, both cell populations are required for learning, suggesting that distinct streams of sensory and task-related information may converge within deep cortical circuits to support adaptive behaviour. Layer 6b (L6b) also receives long-range feedback input^49^, and this could contribute to the deep laminar BOLD signals that we observed. Layer 6b may provide one potential circuit mechanism for top-down attention to influence perception of our memorised images by regulating the activity of L5ET neurons and their interactions with higher-order thalamic circuits^50^. Although our fMRI measurements lack the spatial resolution required to distinguish precise cortical layers, these findings provide an appealing cellular framework for interpreting the differences we observe between superficial and deep cortical signals.

**Fig. 5.**
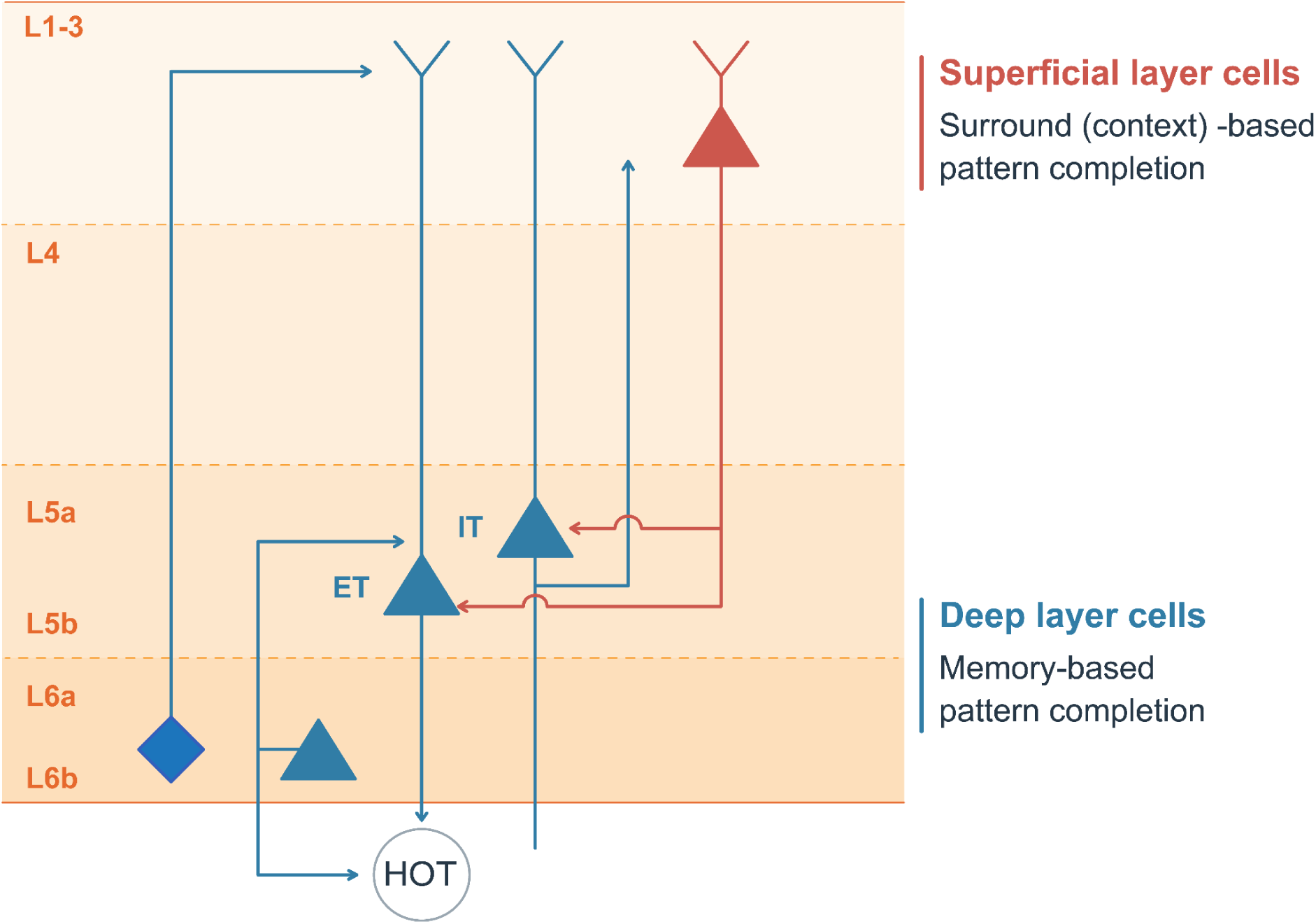
Proposed laminar model for context- and memory-based pattern completion. Schematic diagram illustrating distinct functional roles and connectivity across superficial and deep cortical layers. Superficial layer cells (red) mediate context-based pattern completion, sending output projections to deep-layer sub-populations. Deep-layer cells (blue) serve memory-based pattern completion and comprise distinct subtypes, including layer 5 intratelencephalic (IT) neurons, which primarily receive sensory inputs, and extratelencephalic (ET) neurons, which encode task-relevance signals. These pathways project dynamically to higher-order thalamic (HOT) nuclei and across cortical layers (L1–L3, L4, L5a, L5b, L6a, L6b), providing a cellular framework for layer-dependent signal integration during adaptive behaviour.

Across both ROIs, univariate response amplitudes decreased systematically with repeated exposure, following the pattern Novel > Highly Familiar > Surround completion (repeated single-image). This is consistent with repetition suppression and predictive-processing accounts, in which increasingly predictable stimuli evoke progressively smaller neural responses^12,51,52,53,54^. Importantly, however, category decoding in the feedback ROI emerged only in the Highly Familiar condition. Thus, memory-related improvements in category decoding cannot be explained by stronger neural responses. Instead, repeated experience appears to increase the efficiency and specificity of V1 representations, allowing category-relevant information to be represented more reliably despite lower overall activation. This pattern of results is consistent with sharpening^55,56,57^ or local neuronal down-scaling^54^. The modulation of the deep-to-superficial laminar gradient within the feedback ROI further suggests that familiarity influences not only response magnitude but also the organisation of activity across cortical depths.

The interpretation that V1 representations in the Novel condition are more strongly driven by feedforward visual information, whereas those in the Highly Familiar condition are increasingly shaped by memory-related feedback, is further supported by the representational similarity analyses. Under the Novel condition, V1 activity patterns showed significant correspondence with AlexNet Conv2, indicating that representations were dominated by stimulus-driven visual features, consistent with accounts in which early visual cortex is primarily constrained by low-level image structure^28,30^. In contrast, this correspondence disappeared under the Highly Familiar condition, despite the emergence of reliable category decoding. Together, these findings suggest that repeated experience shifted V1 representations away from low-level feature-based coding and toward memory-constrained representations that supported category-level inference. Importantly, this representational shift occurred alongside reduced overall activity, linking the effect to repetition suppression and predictive-coding accounts in which repeated or expected stimuli evoke smaller responses because they are processed more efficiently^12,51,52,53,54^. Thus, memory-related feedback did not simply amplify V1 responses, but changed the information expressed within V1 while overall signal amplitude decreased.

Overall, our results show that the primary visual cortex expresses distinct forms of predictive information during recognition under occlusion. Prior experience enabled category-level information to emerge within the retinotopic region of V1 corresponding to unseen image content, suggesting that memory-derived predictions can shape early visual representations when local sensory input is incomplete. By contrast, highly informative surround information supported image-specific pattern completion through a different laminar profile, consistent with a more context-driven form of perceptual completion. The dissociation between deep-layer memory-related feedback and superficial-layer surround-based feedback suggests that multiple sources of top-down information converge within V1 through partially distinct pathways. Rather than serving only as a feedforward representation of the visible image, V1 may provide a retinotopic workspace in which memory-based and surround-driven predictions are expressed to support recognition of incomplete visual input.

## Methods

### Participants

41 healthy participants (with normal or corrected-to-normal vision) from the University of Glasgow took part in two experiments (24 females, age range: 18 - 40 years, mean = 24.63 ± 4.77 years), 24 participants for Experiment 1 and 17 participants for Experiment 2; there were no participants who joined both experiments. We paid participants for their participation, and they provided informed written consent. The local ethics committee at the University of Glasgow approved the study (#200220337). In Experiment 1, we excluded six participants: two participants’ data had functional and anatomical alignment problems, one participant did not finish the scanning session, and we were unable to define the region of interest (ROI) for three participants due to scanning artefacts and poor task performance during mapping. Therefore, we report data for 18 participants (nine males; age range: 19-32 years; mean = 25.06 ± 4.36 years). Two of these participants contributed three usable experimental runs; one run was excluded due to excessive head motion or reduced alertness during scanning. In Experiment 2, we excluded two participants for excessive head movement (during anatomical data collection), and one participant for a poorly visible lower part in the image during scanning. Therefore, we report data from 14 participants (eight females; age range: 20-40 years; mean = 25.36 ± 5.54 years).

### Scene Stimuli

We selected 192 real-world scenes containing city and office categories from the Scene Classification Project Upright versus Inverted Scene Database (https://www.bwlab.org/datacode; ^58^). In Experiment 1, we compiled a set of 192 scene stimuli, consisting of 96 city scenes and 96 office scenes, each with a resolution of 800 × 600 pixels. To ensure sufficient variability (and minimise the effect of similarity on memory performance), we replaced highly similar images with additional photographs collected from public image repositories and our own photographs of different city streets and office environments. We converted all images to grayscale and adjusted for global luminance by using Python scripts (OpenCV package). To quantify memory-related effects on feedback signals, we implemented a specific experimental design. To ensure balanced statistical power across conditions, we presented each image an equal number of times (once) inside the scanner. As the “Novel” condition required that stimuli be encountered for the first time during scanning, these images were presented only once. To avoid disproportionate statistical power arising from increased trial numbers in the ‘Familiar’ and ‘Highly Familiar’ conditions, these stimuli were likewise presented only once during scanning, having been pre-exposed during a pre-training phase outside the scanner. As such, in Experiment 1, we used the full set of 192 scene stimuli (96 city scenes and 96 office scenes). In Experiment 2, we compared the effects of memory and surround completion on cortical feedback signals. In Experiment 2, we retained the ‘Novel’ and ‘Highly Familiar’ conditions, randomly selecting 64 images (32 cities and 32 offices) per condition from the 192 scene images, and added two scene images (Fig. 1a, car and market scenes, each shown 32 times) from our previous occlusion paradigm^15^.

### Scene Stimuli features test

#### Structural similarity index model

To reduce the effect of similarity on memory performance, we selected highly dissimilar images using the Structural Similarity Index Model (Python 3.11, scikit-image package). SSIM quantifies image similarity based on luminance, contrast, and structural information ^59^ (Fig. 1 b). We replaced highly similar images with alternatives from the same categories and reassessed image similarity to ensure that the final image set remained dissimilar; similarity values range from 0 to 1 (1 means ‘same’)^59^. Across all stimuli (city and office), the mean similarity was 0.22 ± 0.07, confirming that the stimuli were sufficiently distinct and minimising potential similarity effects on memory performance.

#### Gist model features

To quantify the structural properties of the stimuli and account for low-level visual similarities, we extracted GIST-like image features. We used Python 3.11 (with the OpenCV package; ^60^) to construct a set of Gabor filters to capture spatial-frequency and orientation information. We generated Gabor kernels with four spatial frequencies (0.05, 0.1, 0.2, 0.4 cycles/pixel) and eight orientations uniformly distributed over the interval [0, π], yielding a total of 32 filters. We convolved each grayscale image with these filters, producing 32 corresponding response maps. To capture spatial layout, we divided each response map into a 4 × 4 grid and computed the mean filter response within each grid cell, which yielded 16 values per filter and a total feature vector of 512 dimensions (32 filters × 16 regions), providing a compact representation of the image’s low-level structure. This approach approximates the original GIST descriptor by summarising oriented spatial frequency content across the image in a coarse spatial layout^61^. We computed a representational similarity matrix with a 192 × 192 scale, where the dissimilarity between each pair of images was defined as 1 - Pearson’s correlation coefficient, capturing the representational distinction between city and office images^61,62^. Fig. 1b shows the GIST feature representational dissimilarity matrix (RDM) for the sorted images in the city and office categories. GIST-based dissimilarity values were compared across within-City, within-Office, and City-Office image pairs. Levene’s test indicated heterogeneity of variance (W = 5.44, p = 0.004), so Welch’s one-way ANOVA was used, revealing a significant effect of comparison type [F(2, 9884.08) = 54.65, p < 0.001]. Post hoc comparisons showed that within-City dissimilarity was lower than both City-Office and within-Office dissimilarities (both p < 0.001), whereas City-Office and within-Office dissimilarities did not differ (p = 0.52). Thus, although City images showed somewhat greater within-category similarity, there was no evidence that between-category image pairs were more dissimilar than within-Office pairs based on these low-level GIST features.

#### AlexNet model features

Building on the GIST analyses, which primarily captured low-level scene statistics, we examined higher-order visual representations using AlexNet. Analyses were performed in Python 3.11 using custom scripts that incorporated PyTorch (v2.0), torchvision, SciPy, NumPy, and pandas. Stimulus features were extracted using AlexNet pretrained on the Places365 scene-recognition dataset^33^. To accommodate grayscale images, the pretrained first-layer RGB filters were averaged across colour channels to initialise a single-channel convolutional layer while preserving the pretrained spatial-filter structure. Images were resized from 800 × 600 to 224 × 224 pixels before feature extraction. Features were sampled across the AlexNet hierarchy, from early convolutional layers to fully connected layers. Early layers such as Conv1 predominantly encode lower-level visual properties, including edges, orientations and texture-like information, whereas later layers such as FC7 capture more abstract and category-related representations^29,32,33,61,63^.

For each network layer, feature maps were flattened into one-dimensional vectors, and pairwise dissimilarity was calculated as one minus the Pearson correlation between stimulus feature vectors, yielding a 192 × 192 representational dissimilarity matrix (RDM; ^61,62^). To assess category structure across the AlexNet hierarchy (Supplementary Fig. 5a), dissimilarities were grouped into within-City, within-Office and between-category (City–Office) pairs. Levene’s tests indicated heterogeneity of variance at both Conv1 and FC7 (both ps < 0.01); therefore, group differences were assessed using Welch’s one-way ANOVA followed by post hoc pairwise comparisons. At Conv1, pair type had a significant effect, F(2, 9737.15) = 52.20, p < 0.001, although the absolute differences between pair types were small (0.002–0.007), indicating only weak category structure at this early feature-processing stage. In contrast, FC7 showed a substantially stronger effect of pair type, F(2, 9818.61) = 4586.66, p < 0.001. Between-category dissimilarities were significantly higher than both within-City and within-Office dissimilarities, with between-category differences of 0.147–0.173, indicating robust separation of City and Office representations at this higher-level network stage. Together, these results show that category separation was weak in early visual-feature representations but became substantially more pronounced in higher-level AlexNet representations (Fig. 1b and Supplementary Fig. 5a).

### Experimental conditions

In Experiment 1, we used a paradigm with three familiarity conditions across two scene categories: City and Office. The familiarity conditions were Novel, Familiar and Highly Familiar, corresponding to images presented once, twice or eight times, respectively. To avoid image-specific confounds, the 192 images were randomly assigned to the three familiarity conditions separately for each participant, while maintaining equal numbers of City and Office scenes in each condition. Thus, each participant had an individualised stimulus list in which different images were assigned to the Novel, Familiar and Highly Familiar conditions. This participant-specific list was then used to determine stimulus presentation across sessions and runs. Within each condition, the 64 images (32 City and 32 Office) were divided equally among four experimental runs, with 16 images per run (eight City and eight Office). During pre-training, only Familiar and Highly Familiar images were presented, whereas Novel images were reserved for their first presentation during scanning. During the fMRI session, the same run-wise allocation was used, such that each run contained 16 Novel, 16 Familiar and 16 Highly Familiar images, with all 192 images presented once across the four runs.

Experiment 2 combined the multi-image category-memory paradigm with a single-image occlusion paradigm (two images were repeatedly presented for inducing surround-based completion). This experiment included two familiarity conditions, Novel and Highly Familiar, across the two scene categories, together with two single images selected from a previous study^15^. Every single image was presented 32 times during scanning, matching the number of repetitions within one category-memory condition. Both experiments consisted of three sessions: a pre-training behavioural session, an fMRI scanning session and a post-test behavioural session (Fig. 1a).

### Behavioural study design and procedure

Experimental stimuli were presented using PsychoPy (version 2022.3.4; ^64^) in both the behavioural and fMRI sessions, on a 1920 × 1080 pixel display with a refresh rate of 144 Hz. Before scanning, participants completed a training session in which full images from the Familiar condition were presented once and images from the Highly Familiar condition were presented seven times. The training session consisted of four runs, with 16 Familiar and 16 Highly Familiar images assigned to each run (Fig. 1a). Each run therefore contained 128 trials: 16 Familiar trials and 112 Highly Familiar trials (16 Highly Familiar images × 7 times). Each image was presented for 4 s, followed by a 0.5-s blank interstimulus interval.

Participants indicated whether each image had previously appeared during the training session. They pressed the right arrow on the keyboard for “No, I have not seen this image before” and the left arrow for “Yes, I have seen this image before.” Accordingly, Highly Familiar images required a “No” response on their first presentation and a “Yes” response on the subsequent six presentations, whereas Familiar images always required a “No” response. In Experiment 2, the Familiar images were retained during training as a Pre-New condition to maintain participants’ attention to the recognition task. Within each pre-training run, the 16 Highly Familiar images were organised into seven repetition groups, each containing one independently randomised presentation of all 16 images. The 16 Familiar (‘Pre-New’ in Experiment 2) trials were then randomly interleaved across these groups. This design ensured that “No” responses remained necessary throughout the run and prevented participants from adopting a simple “Yes” response strategy during the later repetition groups.

During the scanning session, participants were also required to press the button to recognise the current image as ‘OLD/NEW’, which means they should press the ‘No’ button for the ‘Novel’ condition and press the ‘Yes’ button for the ‘Familiar’ and ‘Highly Familiar’ conditions because they had already seen the image in the training session. In Experiment 2, the surround completion condition (32 repetitions for every single image) also had the same task of pressing the ‘No’ button as ‘NEW’ for the first time and ‘Yes’ for the rest of the presentation times (2nd to 32nd).

To test memory performance again, we also designed a post-testing behavioural session to be conducted at the end of the scanning session. We randomly selected eight pairs of city and office images from each familiarity condition. We added 16 new ‘Post-New’ images (equal numbers in city and office), which were presented to participants at random within the same recognition task. The duration of the stimulation presentation (4s) was the same as the pre-training session (Fig. 1a).

### Behavioural data analysis

Experiment 1 included 18 participants. Behavioural response data were available for all 18 participants during pre-training and post-testing behavioural sessions, and for 17 participants during the scanning session. Experiment 2 included 14 participants, with behavioural data available from 13 participants during pre-training, 12 during scanning, and all 14 during post-testing. Missing behavioural datasets were caused by technical problems with the MRI-compatible response button box or stimulus-presentation computer; the corresponding fMRI datasets were retained when image acquisition and task presentation were otherwise completed successfully.

#### Response and reaction time analysis

We first calculated the proportion of “yes” responses for each condition across the pre-training, scanning and post-test sessions. “Yes” responses corresponded to trials on which participants judged the image as previously seen, according to their counterbalanced response mapping. Because Novel images were always first presentations, the expected response for ‘Novel’ trials was “no”. Thus, the proportion of “yes” responses to Novel trials provided a measure of false recognition.

In both experiments, first presentations during pre-training were expected to elicit a “no” response, including the first presentation of the Highly Familiar condition in both experiments and the Familiar condition in Experiment 1 (Pre-New condition in Experiment 2). To characterise the emergence of familiarity across repeated exposure, we calculated the proportion of “yes” responses at each presentation of the Highly Familiar images during training. For Experiment 2, we additionally quantified responses to subsequent presentations of the repeated single-image stimuli during scanning. Reaction times were calculated for valid-response trials in each condition (Fig. 3a, b and Supplementary Fig. 10).

Statistical analyses were performed in Python v.3.11 using pandas, NumPy, SciPy, statsmodels, pingouin and scikit-posthocs. For response rates and reaction times, normality was assessed using the Shapiro–Wilk test, and homogeneity of variance was assessed using Levene’s test. When assumptions of normality and equal variance were met, condition effects were tested using one-way ANOVA followed by Tukey’s honestly significant difference tests. When normality was met, but homogeneity of variance was violated, Welch’s ANOVA, followed by Games–Howell post hoc tests, was used. When normality was violated, the Kruskal–Wallis test was followed by Dunn’s tests with a Benjamini–Hochberg correction. All tests were two-tailed, and statistical significance was defined as p < 0.05. Statistical tests were performed for behavioural data from all three sessions in both experiments. Data visualisation was performed using matplotlib and seaborn.

#### Signal detection analysis

Recognition sensitivity during scanning was quantified using signal detection theory for the behavioural task of the scanning session. For each participant and experiment, responses were aggregated across available runs separately for each condition. In both experiments, trials of the Novel condition were treated as the noise condition. In Experiment 1, Familiar and Highly Familiar trials were treated as signal conditions. In Experiment 2, Highly Familiar trials and single-image control trials were treated as signal conditions. For the single-image control condition in Experiment 2, the first presentation of each repeated single-image stimulus was excluded to ensure that this condition reflected responses to repeated presentations of the same occluded image. For each signal condition, hits were defined as “yes” responses to signal trials, whereas false alarms were defined as “yes” responses to ‘Novel’ trials. Hit rates and false-alarm rates were calculated for each participant and condition. To avoid infinite values when response rates were equal to 0 or 1, rates were corrected using a standard log-linear correction. Recognition sensitivity was then calculated as d′, with higher values indicating better discrimination between previously seen and Novel images.

In Experiment 1, d′ was compared between the Familiar and Highly Familiar conditions to assess the effect of prior exposure on recognition sensitivity. In Experiment 2, d′ was compared between the Highly Familiar and Surround Completion (repeated single-image) conditions to test whether memory-based familiarity differed from repeated exposure to the same occluded image. For each comparison, normality of the paired differences was assessed using the Shapiro–Wilk test. Paired-samples t-tests were used when the normality assumption was met; otherwise, Wilcoxon signed-rank tests were applied.

### fMRI experimental study design and procedure

Both fMRI experiments consisted of four experimental runs, each containing forty-eight ‘natural scene’ trials and three types of ‘mapping’ trials. Natural scene trials comprised six conditions (Novel, Familiar, Highly Familiar for both cities and offices), and the three mapping trials (target, border, and surround); each run lasted approximately 12.2 minutes. The main trials comprised a 6-s stimulus presentation, during which stimuli were flashed at 400 ms on/400 ms off (eight presentations per trial) to enhance the signal-to-noise ratio of the fMRI response^27^. Trials were separated by a jittered interstimulus interval of 4–10 s (mean - 7 s). We presented visual stimulation on a projector with a 60 Hz refresh rate. A tilted mirror installed on the head coil allowed participants to view the stimuli during the experiment, using a rear-projection screen (1024 x 768 resolution, 60Hz refresh rate). We partially occluded natural scene images by placing a uniform grey rectangle over the lower-right quadrant of the image (Fig. 1a; ^8,11,15,16,18,65^). We refer to the V1 representation of the occluded lower-right quadrant as the occluded V1 region, or feedback-dominated region, because it did not receive local feedforward information about the underlying scene content (Fig. 1a and Fig. 2). The surrounding visible feedforward region refers to the three-quarters of the scene that are not covered by the lower-right quadrant of the image. At the end of each run, mapping trials were conducted once for each condition (Fig. 1c). Participants were required to fixate on the central cross during scanning, and clear instructions were provided before scanning, with reminders given before each experimental run. We also scanned for a retinotopic mapping run after all experimental runs, which allowed us to localise the boundaries of early visual areas V1-V3. The mapping stimulus was presented using polar-angle high-contrast flickering checkerboards (Fig. 1c). We engaged the participants to focus on the central dot and spot changes in fixation colour.

### MRI acquisition

We collected MRI data at the Imaging Centre of Excellence, Queen Elizabeth University Hospital, University of Glasgow. We measured blood oxygen level-dependent (BOLD) signals using a 7T Siemens Magnetom Terra MRI scanner with the NOVA head coil (32-channel). Functional scanning comprised a CMRR MB (Multiband factor: 2) echo-planar imaging (2D - EPI) sequence [Echo time (TE): 26.3 ms; repetition time (TR): 2000 ms; field of view (FOV): 148 mm; flip angle: 77°; 22 slices; isotropic spatial resolution: 0.8 × 0.8 × 0.8 mm³; GRAPPA acceleration factor: 3; 24 phase-encoding reference lines acquired using FLEET; phase-encoding direction: posterior-to-anterior] to acquire partial brain volumes aligned to maximise coverage of early visual areas (Fig. 1a). We collected fMRI data from four experimental runs (351 volumes per run) for all participants, plus eight top-up scans; an additional run (397 volumes) was used for retinotopic mapping (polar angle), with two top-up scans. A 3D high-resolution anatomical scan [T1 contrast, 3D Magnetisation Prepared Rapid Gradient Echo (MP2RAGE), repetition time (TR): 4680ms, echo time (TE): 2.07ms, the field of view (FOV): 240 mm, spatial resolution: 0.63 x 0.63 x 0.63 mm³, TI1 = 840 ms, TI2 = 2370 ms, 256 volumes] was also performed in the middle of the scanning session. We collected two top-up scans from AP (anterior to posterior) and PA (posterior to anterior) directions [echo time (TE): 26.3ms, repetition time (TR): 2000ms, the field of view (FOV): 148 mm, flip angle: 77 °, 22 slices, spatial resolution: 0.8 x 0.8 x 0.8 mm³, 5 volumes] before each experimental run and retinotopic mapping run for the distortion correction.

### Functional and Anatomical Data Preprocessing

We performed distortion correction of the 7T functional data using FSL^66^. For each run, we acquired pairs of spin-echo EPI images with reversed phase-encoding directions (posterior–anterior and anterior–posterior) and used these to estimate susceptibility-induced field inhomogeneities, which we subsequently applied to correct geometric distortions in the functional data. We further used BrainVoyager (version 23.2; ^67^) for pre-processing functional data. The preprocessing involves three steps: 3D motion correction (using trilinear/sinc interpolation), slice scan time correction (adjusted by cubic spline interpolation), and a GLM-Fourier high-pass filter (a cycle of 5 sines/cosines for four experimental runs and a cycle of 7 sines/consines for the retinotopy run) for temporal filtering. To improve the alignment between functional and anatomical data, we used a boundary-based registration function as a fine-tuning step, allowing rough alignment checks by visual inspection. We also calculated the between-run alignment using Pearson’s r correlation to ensure all functional runs were well aligned (r > 0.9). The transformation was applied to correct runs with a lower alignment value (r < 0.9) until the standard was reached. We processed the anatomical data using a denoising function in BrainVoyager (23.2) and subsequently resampled to a resolution of 0.4 mm following brain extraction. We first manually corrected the white matter of the native brain to improve the boundary-based co-registration. After transforming anatomical images to standard anterior commissure–posterior commissure (ACPC) brain space, we also manually corrected the white matter and grey matter segmentation by using ITK-SNAP (version 4.1.0; ^68^) to ensure accurate delineation of the white-matter/grey-matter and pial boundaries, which were used to estimate cortical depth and assign voxels to cortical layers. We focused the manual segmentation on two defined regions in V1: one that responded to the visible scene surround (for ease, we name the region as the “feedforward” area, though it also receives feedback signals), and one that responded to the occluded scene region (our “feedback” area of V1, Fig. 1c). This approach ensured that the grey matter segmentation did not involve any parts of the larger veins or skull. After GM/WM segmentation, we reconstructed the surface of the occipital lobe for the following ROI definition.

### Mapping the V1 regions of interest

The following mapping procedure localised the masked image region, allowing us to identify the retinotopic regions of the V1, V2, and V3 cortices that respond to the occluded quadrant. The full stimulus image spanned 19.5 × 14.7° of visual angle, with the grey occluder occupying ∼9 × 7° in the lower-right quadrant. We presented a centralised fixation with a white cross (0.1 x 0.1°) in the centre of the scene images. During the mapping trials, participants viewed high-contrast flickering checkerboards (4 Hz) at one of three visual locations: one at the lower-right image quadrant (Target mapping), one over the image region not containing the Target (Surround mapping), and one at the border between the Target and Surround (Border mapping; Fig 1c). The Target and Surround stimuli matched the experimental stimuli’s size, occupying a region of 9 x 7° visual angle and a region of 19.5 x 14.7° visual angle, while the border regions mapped between the target and surround (approximately 0.5 diagonally from fixation; Fig. 1c) to define the cortical representation of the occluded area and non-occluded area in the primary visual cortex. To define boundaries between V1/V2 and V2/V3, we applied a Fourier transform to each voxel’s fMRI time series from the polar-angle mapping runs, thereby computing both the amplitude and phase at the stimulation frequency. The resulting phase angles were colour-encoded and projected onto the inflated cortical surface. Each colour represented an F-ratio, defined as the squared amplitude at the stimulation frequency divided by the mean squared amplitude across all other frequencies. Based on the colour encoding, we then manually identified the boundary separating V1 from V2 on the cortical surface for each participant (Fig. 1b and Supplementary Fig. 7). We further defined the V1 occluded area in the left hemisphere using a GLM contrast [Target - surround - border] to identify the activated area (Fig. 1c and Supplementary Fig. 7) for each participant. As a stimulated control region, we defined right-hemisphere V1, which represented the contralateral visible portion of the scene and therefore received direct feedforward input from non-occluded image content. We then used MATLAB to exclude the voxels with a mean signal intensity of ≤100 in any run. The remaining voxels were functionally selected using independent mapping data. For the V1 ROIs, voxels showing a target-versus-border response of t ≥ 1 were selected.

### Cortical depth-layer segmentation

Cortical depth–dependent analyses were performed using the high-resolution cortical depth grid-sampling procedure implemented in BrainVoyager. Following white matter and grey matter segmentation and surface reconstruction, cortical thickness was estimated for each participant while accounting for local cortical curvature. These estimates were used to generate subject-specific cortical depth grids across the whole brain. An equivolume approach was applied to subdivide the grey matter ribbon into six depth compartments, ensuring approximately equal volume across depths despite variations in cortical folding^69,70^. Depth grids were then intersected with predefined regions of interest (ROIs) in functional space to obtain depth-resolved voxel subsets for each ROI. The resulting cortical depth spanned the cortical ribbon from the white matter–grey matter boundary to the grey matter–cerebrospinal fluid interface and was defined at relative cortical depths of 0.90, 0.74, 0.58, 0.42, 0.26 and 0.10. Depth values were ordered from the white matter boundary towards the pial surface, with 0.90 corresponding to the deepest compartment and 0.10 corresponding to the most superficial compartment (Fig. 1c). These depth bins reflect relative positions within the cortical ribbon and should not be interpreted as directly corresponding to histologically defined cortical layers.

To examine whether voxel counts differed across cortical depths, we fitted a linear mixed-effects model (R v4.4.2; lme4) with cortical depth and ROI (V1 feedforward and feedback) as fixed effects and participant as a random intercept. The model was specified as:

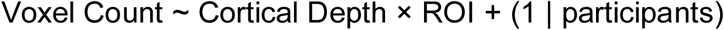

No significant effect of cortical layer was observed [F(1,349) = 0.08, p = 0.78], nor any ROI-specific trends [feedback: t(349) = 0.63, p = 0.53; feedforward: t(349) = −1.04, p = 0.53]. However, voxel counts were significantly higher in feedforward compared to feedback regions [F(1,349) = 910.02, p < 0.001; Supplementary Fig. 8].

### Univariate and Multivoxel Pattern Classification Analysis

Single-trial beta estimates (z-score; code from MATLAB) were extracted using a least-squares-all general linear model (LSA-GLM), and the resulting beta estimates were entered into a linear support vector machine classifier implemented in MATLAB using the LIBSVM toolbox^71^ to decode image-category information from selected voxels within the V1 feedback and feedforward regions. Classification performance was evaluated against the chance level (0.50). Cross-validation was implemented using a leave-one-run-out procedure, in which the classifier was trained on three runs and tested on the remaining run, yielding four classification accuracies per condition for each participant. Decoding accuracy was calculated as the average across validation folds.

To assess memory-related influences on category representations, category decoding (City vs Office) was performed separately for each familiarity condition [Novel, Familiar (Experiment 1 only), and Highly Familiar]. In addition, memory decoding (Novel vs Highly Familiar) was performed in both the V1 feedback region and the feedforward regions. In Experiment 2, decoding of the two repeatedly presented images (Image 1 vs Image 2) was conducted to determine whether the findings replicated those reported by Muckli et al.^15^. To examine laminar differences, the same classification procedure was applied separately to each of the six cortical depths within the V1 feedback and feedforward regions.

Statistical significance was assessed using bootstrap resampling, as classification accuracies are bounded and may not satisfy the normality assumption, especially in small-sample settings. For each condition and cortical depth, 100,000 bootstrap samples were generated by resampling participants with replacement and computing the mean classification accuracy. One-sided bootstrap p values were estimated as the proportion of bootstrap mean accuracies at or below chance level (0.50). 95% confidence intervals were defined by the 2.5th and 97.5th percentiles of the bootstrap distribution. Multiple comparisons were controlled using the false discovery rate (Benjamini–Hochberg procedure; α = 0.05). For each experiment, corrections were applied separately to the feedback and feedforward ROI analyses. Within each ROI, correction was performed across 24 comparisons, corresponding to four conditions tested across six cortical depths. In Experiment 1, the four conditions comprised categorical–Novel, categorical–Familiar, categorical–Highly Familiar, and Novel– Highly Familiar decoding. In Experiment 2, the Familiar–categorical condition was replaced by the surround completion condition (image 1 vs image 2).

To examine condition-related differences in overall BOLD response amplitude, we conducted a complementary univariate analysis using the same single-trial beta estimates. For each trial, beta values were averaged across all selected voxels within the V1 feedback and feedforward ROIs. For laminar analyses, averaging was performed separately within each cortical depth (see Supplementary Fig. 1–4). Trial-wise mean beta values were subsequently averaged across trials belonging to the same condition, yielding one mean beta estimate per participant, condition, ROI, and cortical depth.

### Linear Mixed-Effects Model

In a second-level analysis of layer fMRI MVPA results, we used R 4.4.2 with the nlme package to fit linear mixed-effects (LME) models for several comparisons. The utilisation of the model improves performance in multiple comparisons by parameterising effects across layers and experiments. The framework enhances statistical power by leveraging the accumulation of evidence across conditions. Linear mixed-effects models also allowed us to calculate statistical differences in the same experimental condition across two experiments. To estimate the model, we first extracted the decoding accuracy results from all cortical depths, ROIs, and experimental conditions across the two experiments, for example, effects included ROIs (Feedback vs. Feedforward), cortical depth, experiment (Experiment 1 and Experiment 2), experimental condition (Novel-categorical, Highly Familiar-categorical, and Novel vs Highly Familiar), as well as the interactions between experiment and cortical depth, between experimental condition and cortical depth, and the three-way interaction of experiment, experimental conditions, and cortical depths. The model was specified as:

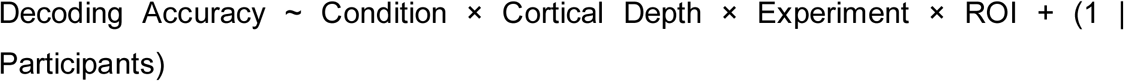

Mean BOLD signal changes (beta estimates) were also analysed using a linear mixed-effects model, with Condition, Layer, ROI, and their interactions as fixed effects, and participants as a random intercept. Follow-up analyses were conducted within the same statistical framework to characterise significant effects across conditions, layers, and ROIs.

### fMRI and AlexNet RDMs Correlation

To further explore the specific feature representations in V1, we performed RDM correlations between fMRI and AlexNet^31,62^. Numerical analyses were performed in Python using NumPy, pandas and SciPy. The fMRI RDMs were computed from voxel-wise beta weights (same as multivoxel-pattern analysis) within the V1 layers of the feedforward and feedback regions, extracted from GLM estimates for each familiarity condition. For each participant, data were sorted by image category (City, Office) and run (1-4), and pairwise dissimilarities between image categories were calculated as 1 - Pearson’s r (Fig. 4a). To ensure a direct comparison, AlexNet RDMs were constructed using image sets matched to the category-run and familiarity conditions in the fMRI experiment. The AlexNet features from each layer were flattened, and pairwise dissimilarities were calculated as 1 - Pearson’s r to generate model RDMs for each condition. Separate RDMs were computed for the Novel and Highly Familiar conditions in both the fMRI and AlexNet conditions.

Image presentation order was randomised within each fMRI run, and the trial-wise randomisation log linking individual image identities to fMRI beta estimates was not recorded. However, image allocation to runs was fixed before scanning, with 16 images assigned to each run. Although image-level pairing was unavailable, the category-by-run structure remained defined by the experimental design. Trial-wise RDMs (64 × 64) for each familiarity condition were reduced to category-by-run RDMs (8 × 8) by averaging dissimilarities within each predefined category × run block (Fig. 4a). In this procedure, correlation distances were averaged across all trial pairs within each run-category-wise block (City runs 1-4; Office runs 1-4), yielding an 8 × 8 RDM (6 × 6 RDM for data from participants with three useable run) in which each cell reflects the mean dissimilarity between category-run blocks (Fig. 4a). AlexNet RDMs were processed with the same reduction to ensure direct comparability. We then performed Spearman rank correlations between each pair of laminar AlexNet and fMRI RDMs by extracting the lower-triangular elements (excluding the diagonal) from each matrix and flattening them into vectors for correlation analysis. To assess statistical significance, correlation values across participants were subjected to a one-sample right-tailed t-test against zero to test whether the correlations were significantly greater than chance. Resulting p-values were corrected for multiple comparisons across 2 ROIs (V1 feedforward and feedback), six cortical depths, seven AlexNet layers and two conditions (Novel and Highly Familiar) by using the Benjamini-Hochberg false discovery rate (FDR) procedure.

## Acknowledgements

This project has received funding from the Medical Research Council (MRC: [UKRI532] ‘Cortical layer-specific imaging of context-dependent cognitive processing’ to LFM and LSP), Biotechnology and Biological Sciences Research Council (BBSRC: BB/V010956/1 ‘Layer-specific cortical feedback’ to LFM with LSP) and the European Union’s Horizon 2020 Framework Programme for Research and Innovation under the Specific Grant Agreement No 945539 (Human Brain Project SGA3) to LFM. We thank F. Crabbe and N. Nothnagel for help in acquiring fMRI data and M. Svanera, X. Yan, J. Ding, Y. Gao, H. Nie, YX, Gao, X. Shao, P. Liu, and Y. Lu for their insightful discussions and support. For the purpose of open access, the author(s) have applied a Creative Commons Attribution (CC BY) licence to any Author Accepted Manuscript version arising from this submission.

## Author information

### Author notes

These authors contributed equally: Lucy S. Petro and Lars F. Muckli.

### Contributions

Conceptualisation - ZZ, CA, MW, CNL, MEL, LSP, LFM. Methodology - ZZ, CA, LSP, LFM. Software - ZZ. Validation - ZZ, CA, LSP, LFM. Formal analysis - ZZ. Investigation - ZZ, YH, YL. Resources - ZZ, CA, ATP. Data curation - ZZ. Writing - original draft - ZZ, LSP, LFM, CA. Writing - review and editing - ZZ, CA, MW, CNL, MEL, LSP, LFM. Visualisation - ZZ. Supervision- LSP, LFM, CA. Project administration - ZZ, LSP, LFM. Funding acquisition - LSP, LFM.

### Corresponding authors

Correspondence to Clement Abbatecola, Lars F. Muckli and Lucy S. Petro.

## Ethics declarations

The authors declare no competing interests.

## Data availability

Raw MRI data from human participants are not publicly available due to data protection requirements but may be shared upon reasonable request following completion of a formal data-sharing agreement and submission of a project outline. Additional information required to reanalyse the data reported in this paper is available from the authors upon reasonable request. Source data underlying the plots and analyses, together with the analysis code, are publicly available at: https://github.com/ZZRdpgyg/7T-M-C-project.

## Code availability

The analysis and plot code is publicly available via GitHub: statistical and model analysis https://github.com/ZZRdpgyg/7T-M-C-project and MVPA https://github.com/Muckli-lab/MVP-analysis-tool-box

## Supplementary Information

Please check on SupplementaryInformation.pdf

## Notes

### Competing Interest Statement

The authors have declared no competing interest.

### Summary of Updates

Author affiliations updated Added line number

## References

1. Squire, L. R., Wixted, J. T. & Clark, R. E. Recognition memory and the medial temporal lobe: a new perspective. Nat. Rev. Neurosci. 8, 872–883 (2007). 10.1038/nrn2154

2. Eichenbaum, H., Yonelinas, A. P. & Ranganath, C. The medial temporal lobe and recognition memory. Annu. Rev. Neurosci. 30, 123–152 (2007). 10.1146/annurev.neuro.30.051606.094328

3. DiCarlo, J. J., Zoccolan, D. & Rust, N. C. How does the brain solve visual object recognition? Neuron 73, 415–434 (2012). 10.1016/j.neuron.2012.01.010

4. Wandell, B. A., Dumoulin, S. O. & Brewer, A. A. Visual field maps in human cortex. Neuron 56, 366–383 (2007). 10.1016/j.neuron.2007.10.012

5. Muckli, L. & Petro, L. S. Network interactions: non-geniculate input to V1. Curr. Opin. Neurobiol. 23, 195–201 (2013). 10.1016/j.conb.2013.01.020

6. Gilbert, C. D. & Li, W. Top-down influences on visual processing. Nat. Rev. Neurosci. 14, 350–363 (2013). 10.1038/nrn3476

7. van Kerkoerle, T., Self, M. W. & Roelfsema, P. R. Layer-specificity in the effects of attention and working memory on activity in primary visual cortex. Nat. Commun. 8, 13804 (2017). 10.1038/ncomms13804

8. Revina, Y., Petro, L. S. & Muckli, L. Cortical feedback signals generalise across different spatial frequencies of feedforward inputs. NeuroImage 180, 280–290 (2018). 10.1016/j.neuroimage.2017.09.047

9. Larkum, M. E., Petro, L. S., Sachdev, R. N. S. & Muckli, L. A perspective on cortical layering and layer-spanning neuronal elements. Front. Neuroanat. 12, 56 (2018). 10.3389/fnana.2018.00056

10. Vezoli, J. et al. Cortical hierarchy, dual counterstream architecture and the importance of top-down generative networks. NeuroImage 225, 117479 (2021). 10.1016/j.neuroimage.2020.117479

11. Smith, F. W. & Muckli, L. Nonstimulated early visual areas carry information about surrounding context. Proc. Natl Acad. Sci. USA 107, 20099–20103 (2010). 10.1073/pnas.1000233107

12. Kok, P., Jehee, J. F. M. & de Lange, F. P. Less is more: expectation sharpens representations in the primary visual cortex. Neuron 75, 265–270 (2012). 10.1016/j.neuron.2012.04.034

13. Vetter, P., Smith, F. W. & Muckli, L. Decoding sound and imagery content in early visual cortex. Curr. Biol. 24, 1256–1262 (2014). 10.1016/j.cub.2014.04.020

14. Bosch, S. E., Jehee, J. F. M., Fernández, G. & Doeller, C. F. Reinstatement of associative memories in early visual cortex is signaled by the hippocampus. J. Neurosci. 34, 7493–7500 (2014). 10.1523/jneurosci.0805-14.2014

15. Muckli, L. et al. Contextual feedback to superficial layers of V1. Curr. Biol. 25, 2690–2695 (2015). 10.1016/j.cub.2015.08.057

16. Morgan, A. T., Petro, L. S. & Muckli, L. Scene representations conveyed by cortical feedback to early visual cortex can be described by line drawings. J. Neurosci. 39, 9410–9423 (2019). 10.1523/jneurosci.0852-19.2019

17. Hayden, D. J. et al. Electrophysiological signatures of visual recognition memory across all layers of mouse V1. J. Neurosci. 43, 7307–7321 (2023). 10.1523/jneurosci.0090-23.2023

18. Petro, L. S., Smith, F. W., Abbatecola, C. & Muckli, L. The spatial precision of contextual feedback signals in human V1. Biology 12, 1022 (2023). 10.3390/biology12071022

19. Ortiz-Tudela, J. et al. Concurrent contextual and time-distant mnemonic information co-exist as feedback in the human visual cortex. NeuroImage 265, 119778 (2023). 10.1016/j.neuroimage.2022.119778

20. Bergmann, J. et al. Cortical depth profiles in primary visual cortex for illusory and imaginary experiences. Nat. Commun. 15, 1002 (2024). 10.1038/s41467-024-45065-w

21. Thomas, E. R. et al. Predictions and errors are distinctly represented across V1 layers. Curr. Biol. 34, 2265–2271.e4 (2024). 10.1016/j.cub.2024.04.036

22. Lazarova, Y. Y., Huang, Y., Muckli, L. F. & Petro, L. S. Perceptual priors update contextual feedback processing in V1. Top. Cogn. Sci. 10.1111/tops.70018 (2025).

23. Hubel, D. H. & Wiesel, T. N. Receptive fields and functional architecture of monkey striate cortex. J. Physiol. 195, 215–243 (1968). 10.1113/jphysiol.1968.sp008455

24. Freeman, J., Brouwer, G. J., Heeger, D. J. & Merriam, E. P. Orientation decoding depends on maps, not columns. J. Neurosci. 31, 4792–4804 (2011). 10.1523/jneurosci.5160-10.2011

25. Haynes, J.-D. & Rees, G. Predicting the orientation of invisible stimuli from activity in human primary visual cortex. Nat. Neurosci. 8, 686–691 (2005). 10.1038/nn1445

26. Kamitani, Y. & Tong, F. Decoding the visual and subjective contents of the human brain. Nat. Neurosci. 8, 679–685 (2005). 10.1038/nn1444

27. Kay, K. N., Naselaris, T., Prenger, R. J. & Gallant, J. L. Identifying natural images from human brain activity. Nature 452, 352–355 (2008). 10.1038/nature06713

28. Khaligh-Razavi, S.-M. & Kriegeskorte, N. Deep supervised, but not unsupervised, models may explain IT cortical representation. PLoS Comput. Biol. 10, e1003915 (2014). 10.1371/journal.pcbi.1003915

29. Güçlü, U. & van Gerven, M. A. J. Deep neural networks reveal a gradient in the complexity of neural representations across the ventral stream. J. Neurosci. 35, 10005–10014 (2015). 10.1523/jneurosci.5023-14.2015

30. Yamins, D. L. K. & DiCarlo, J. J. Using goal-driven deep learning models to understand sensory cortex. Nat. Neurosci. 19, 356–365 (2016). 10.1038/nn.4244

31. Cichy, R. M., Khosla, A., Pantazis, D., Torralba, A. & Oliva, A. Comparison of deep neural networks to spatio-temporal cortical dynamics of human visual object recognition reveals hierarchical correspondence. Sci. Rep. 6, 27755 (2016). 10.1038/srep27755

32. Krizhevsky, A., Sutskever, I. & Hinton, G. E. ImageNet classification with deep convolutional neural networks. In Advances in Neural Information Processing Systems 25 (eds Pereira, F., Burges, C. J. C., Bottou, L. & Weinberger, K. Q.) 1097–1105 (Curran Associates, 2012). <u>Source</u>

33. Zhou, B., Lapedriza, A., Khosla, A., Oliva, A. & Torralba, A. Places: a 10 million image database for scene recognition. IEEE Trans. Pattern Anal. Mach. Intell. 40, 1452–1464 (2018). 10.1109/tpami.2017.2723009

34. Wagatsuma, N., Hidaka, A. & Tamura, H. Analysis based on neural representation of natural object surfaces to elucidate the mechanisms of a trained AlexNet model. Front. Comput. Neurosci. 16, 979258 (2022). 10.3389/fncom.2022.979258

35. Berezovskii, V. K., Nassi, J. J. & Born, R. T. Segregation of feedforward and feedback projections in mouse visual cortex. J. Comp. Neurol. 519, 3672–3683 (2011). 10.1002/cne.22675

36. Markov, N. T. et al. Anatomy of hierarchy: feedforward and feedback pathways in macaque visual cortex. J. Comp. Neurol. 522, 225–259 (2014). 10.1002/cne.23458

37. Finnie, P. S. B., Komorowski, R. W. & Bear, M. F. The spatiotemporal organization of experience dictates hippocampal involvement in primary visual cortical plasticity. Curr. Biol. 31, 3996–4008.e6 (2021). 10.1016/j.cub.2021.06.079

38. Badwal, M. W., Bergmann, J., Roth, J. H. R., Doeller, C. F. & Hebart, M. N. The scope and limits of fine-grained image and category information in the ventral visual pathway. J. Neurosci. 45, e0936242024 (2025). 10.1523/jneurosci.0936-24.2024

39. Rao, R. P. N. & Ballard, D. H. Predictive coding in the visual cortex: a functional interpretation of some extra-classical receptive-field effects. Nat. Neurosci. 2, 79– 87 (1999). 10.1038/4580

40. Petro, L. S., Vizioli, L. & Muckli, L. Contributions of cortical feedback to sensory processing in primary visual cortex. Front. Psychol. 5, 1223 (2014). 10.3389/fpsyg.2014.01223

41. Marr, D. Simple memory: a theory for archicortex. Philos. Trans. R. Soc. Lond. B Biol. Sci. 262, 23–81 (1971). 10.1098/rstb.1971.0078

42. McClelland, J. L., McNaughton, B. L. & O’Reilly, R. C. Why there are complementary learning systems in the hippocampus and neocortex: insights from the successes and failures of connectionist models of learning and memory. Psychol. Rev. 102, 419–457 (1995). 10.1037/0033-295x.102.3.419

43. Staresina, B. P. & Wimber, M. A neural chronometry of memory recall. Trends Cogn. Sci. 23, 1071–1085 (2019). 10.1016/j.tics.2019.09.011

44. Theves, S., Grande, X., Düzel, E. & Doeller, C. F. Pattern completion and the medial temporal lobe memory system. In The Oxford Handbook of Human Memory (eds Kahana, M. J. & Wagner, A. D.) 988–1016 (Oxford Univ. Press, 2024). 10.1093/oxfordhb/9780190917982.013.34

45. Hindy, N. C., Avery, E. W. & Turk-Browne, N. B. Hippocampal-neocortical interactions sharpen over time for predictive actions. Nat. Commun. 10, 3989 (2019). 10.1038/s41467-019-12016-9

46. Kaefer, K., Stella, F., McNaughton, B. L. & Battaglia, F. P. Replay, the default mode network and the cascaded memory systems model. Nat. Rev. Neurosci. 23, 628–640 (2022). 10.1038/s41583-022-00620-6

47. Jordan, R. & Keller, G. B. Opposing influence of top-down and bottom-up input on excitatory layer 2/3 neurons in mouse primary visual cortex. Neuron 108, 1194– 1206 (2020). 10.1016/j.neuron.2020.09.024

48. Moberg, S. et al. Distinct roles of cortical layer 5 subtypes in associative learning. Nat. Commun. 17, 2648 (2026). 10.1038/s41467-026-68307-5

49. Zolnik, T. A. et al. Layer 6b is driven by intracortical long-range projection neurons. Cell Rep. 30, 3492–3505.e5 (2020). 10.1016/j.celrep.2020.02.044

50. Zolnik, T. A., Eickholt, B. J., Molnár, Z. & Larkum, M. E. The layer 6b theory of attention. Neuron 114, 565–582 (2026). 10.1016/j.neuron.2025.11.024

51. Grill-Spector, K., Henson, R. & Martin, A. Repetition and the brain: neural models of stimulus-specific effects. Trends Cogn. Sci. 10, 14–23 (2006). 10.1016/j.tics.2005.11.006

52. Gotts, S. J., Chow, C. C. & Martin, A. Repetition priming and repetition suppression: a case for enhanced efficiency through neural synchronization. Cogn. Neurosci. 3, 227–237 (2012). 10.1080/17588928.2012.670617

53. Auksztulewicz, R. & Friston, K. J. Repetition suppression and its contextual determinants in predictive coding. Cortex 80, 125–140 (2016). 10.1016/j.cortex.2015.11.024

54. Alink, A., Abdulrahman, H. & Henson, R. N. Forward models demonstrate that repetition suppression is best modelled by local neural scaling. Nat. Commun. 9, 3854 (2018). 10.1038/s41467-018-05957-0

55. Barron, H. C., Garvert, M. M. & Behrens, T. E. J. Repetition suppression: a means to index neural representations using BOLD? Philos. Trans. R. Soc. Lond. B Biol. Sci. 371, 20150355 (2016). 10.1098/rstb.2015.0355

56. Cui, L. et al. Stimulus repetition induces a two-stage learning process in the primary visual cortex. J. Neurosci. 45, e1788242025 (2025). 10.1523/jneurosci.1788-24.2025

57. Seignette, K. et al. Opposing plasticity mechanisms in single neurons shape visual saliency assignment. bioRxiv (2024). 10.1101/2024.06.10.598181

58. Walther, D. B., Chai, B., Caddigan, E., Beck, D. M. & Fei-Fei, L. Simple line drawings suffice for functional MRI decoding of natural scene categories. Proc. Natl Acad. Sci. USA 108, 9661–9666 (2011). 10.1073/pnas.1015666108

59. Wang, Z., Bovik, A. C., Sheikh, H. R. & Simoncelli, E. P. Image quality assessment: from error visibility to structural similarity. IEEE Trans. Image Process. 13, 600– 612 (2004). 10.1109/tip.2003.819861

60. Bradski, G. The OpenCV library. Dr. Dobb’s J. Softw. Tools 25, 120–125 (2000). Source

61. Lahner, B., Mohsenzadeh, Y., Mullin, C. & Oliva, A. Visual perception of highly memorable images is mediated by a distributed network of ventral visual regions that enable a late memorability response. PLoS Biol. 22, e3002564 (2024). 10.1371/journal.pbio.3002564

62. Kriegeskorte, N., Mur, M. & Bandettini, P. A. Representational similarity analysis— connecting the branches of systems neuroscience. Front. Syst. Neurosci. 2, 4 (2008). 10.3389/neuro.06.004.2008

63. Yamins, D. L. K., et al. Performance-optimized hierarchical models predict neural responses in higher visual cortex. Proc. Natl Acad. Sci. USA 111, 8619–8624 (2014). 10.1073/pnas.1403112111

64. Peirce, J. W. et al. PsychoPy2: experiments in behavior made easy. Behav. Res. Methods 51, 195–203 (2019). 10.3758/s13428-018-01193-y

65. Papale, P. et al. The representation of occluded image regions in area V1 of monkeys and humans. Curr. Biol. 33, 3865–3871.e3 (2023). 10.1016/j.cub.2023.08.010

66. Jenkinson, M., Beckmann, C. F., Behrens, T. E. J., Woolrich, M. W. & Smith, S. M. FSL. NeuroImage 62, 782–790 (2012). 10.1016/j.neuroimage.2011.09.015

67. Goebel, R. BrainVoyager—past, present, future. NeuroImage 62, 748–756 (2012). 10.1016/j.neuroimage.2012.01.083

68. Yushkevich, P. A. et al. User-guided 3D active contour segmentation of anatomical structures: significantly improved efficiency and reliability. NeuroImage 31, 1116–1128 (2006). 10.1016/j.neuroimage.2006.01.015

69. Wähnert, M. D. et al. Anatomically motivated modeling of cortical laminae. NeuroImage 93, 210–220 (2014). 10.1016/j.neuroimage.2013.03.078

70. Kemper, V. G., De Martino, F., Emmerling, T. C., Yacoub, E. & Goebel, R. High resolution data analysis strategies for mesoscale human functional MRI at 7 and 9.4T. NeuroImage 164, 48–58 (2018). 10.1016/j.neuroimage.2017.03.058

71. Chang, C.-C. & Lin, C.-J. LIBSVM: a library for support vector machines. ACM Trans. Intell. Syst. Technol. 2, 27 (2011). 10.1145/1961189.1961199

