## Supplementary information for "Memory-based and surround-based pattern completion show distinct laminar profiles in human V1"

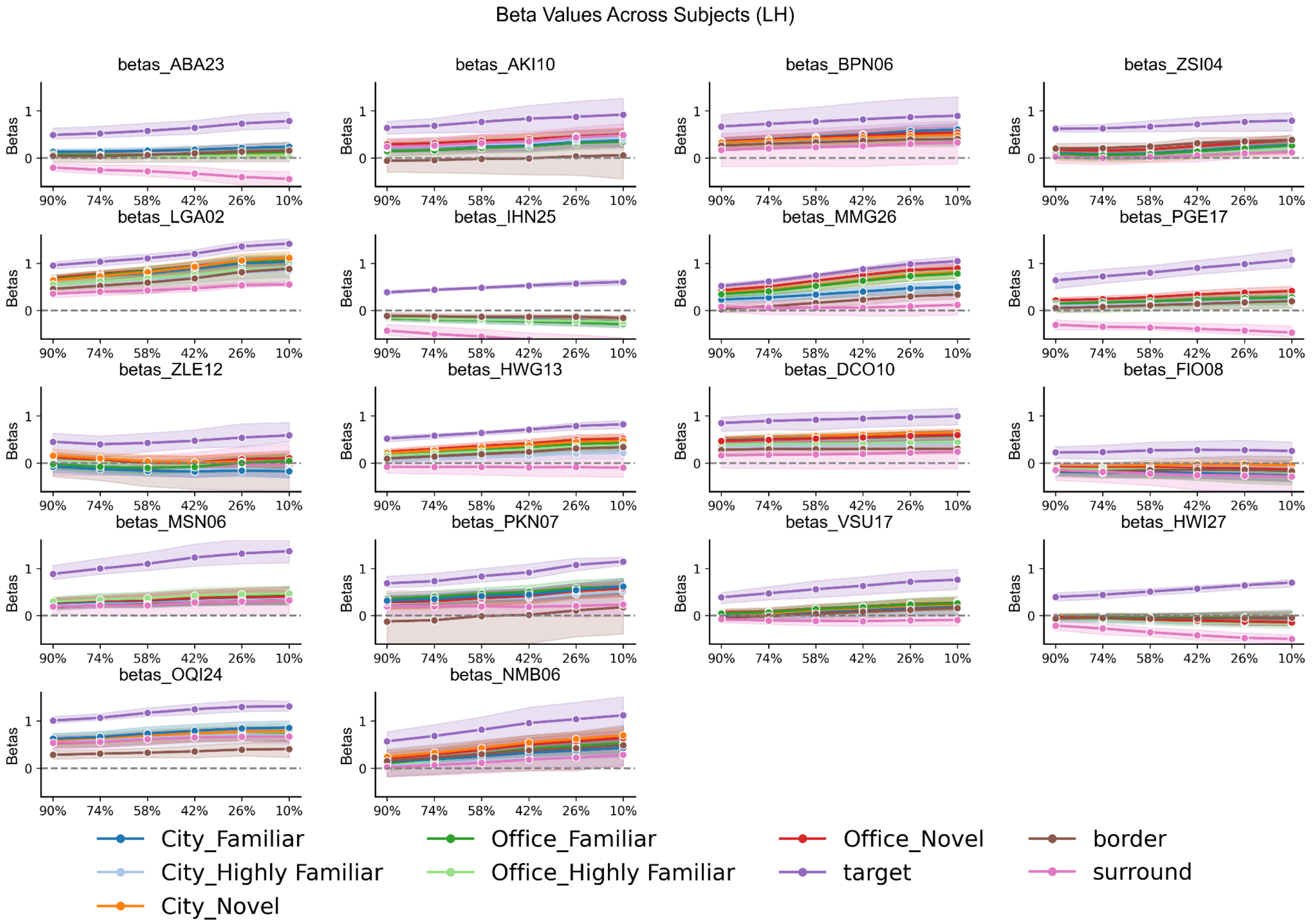


**Supplementary Fig. 1. Individual univariate BOLD betas across all layers in the V1 feedback area in Experiment 1 (N = 18).** Shown are mean beta estimates for the three mapping conditions (surround, border and target) and the six experimental conditions: City Novel, City Familiar, City Highly Familiar, Office Novel, Office Familiar and Office Highly Familiar.

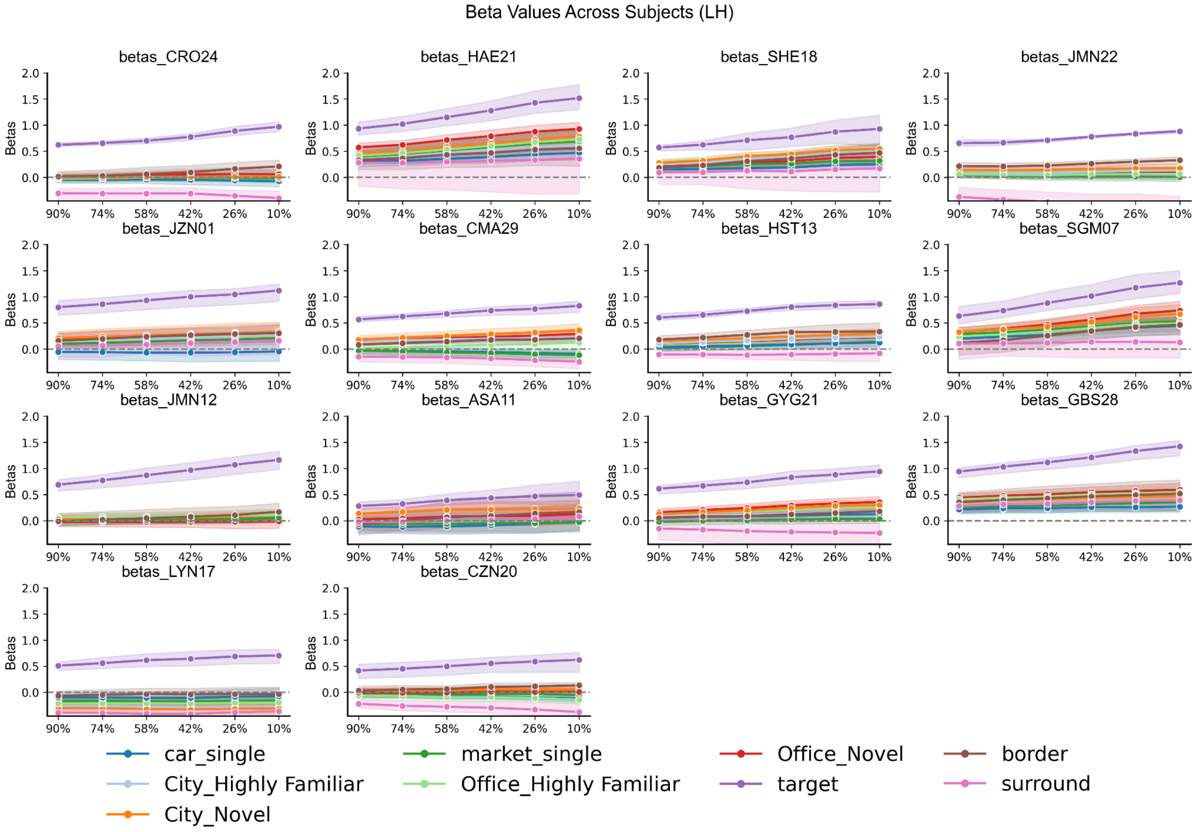


**Supplementary Fig. 2. Individual univariate BOLD betas across all layers in the V1 feedback area in Experiment 2 (N = 14).** Shown are mean beta estimates for the three mapping conditions (surround, border and target) and the six experimental conditions: City Novel, car-single (surround completion image 1), City Highly Familiar, Office Novel, market-single (surround completion image 2) and Office Highly Familiar.

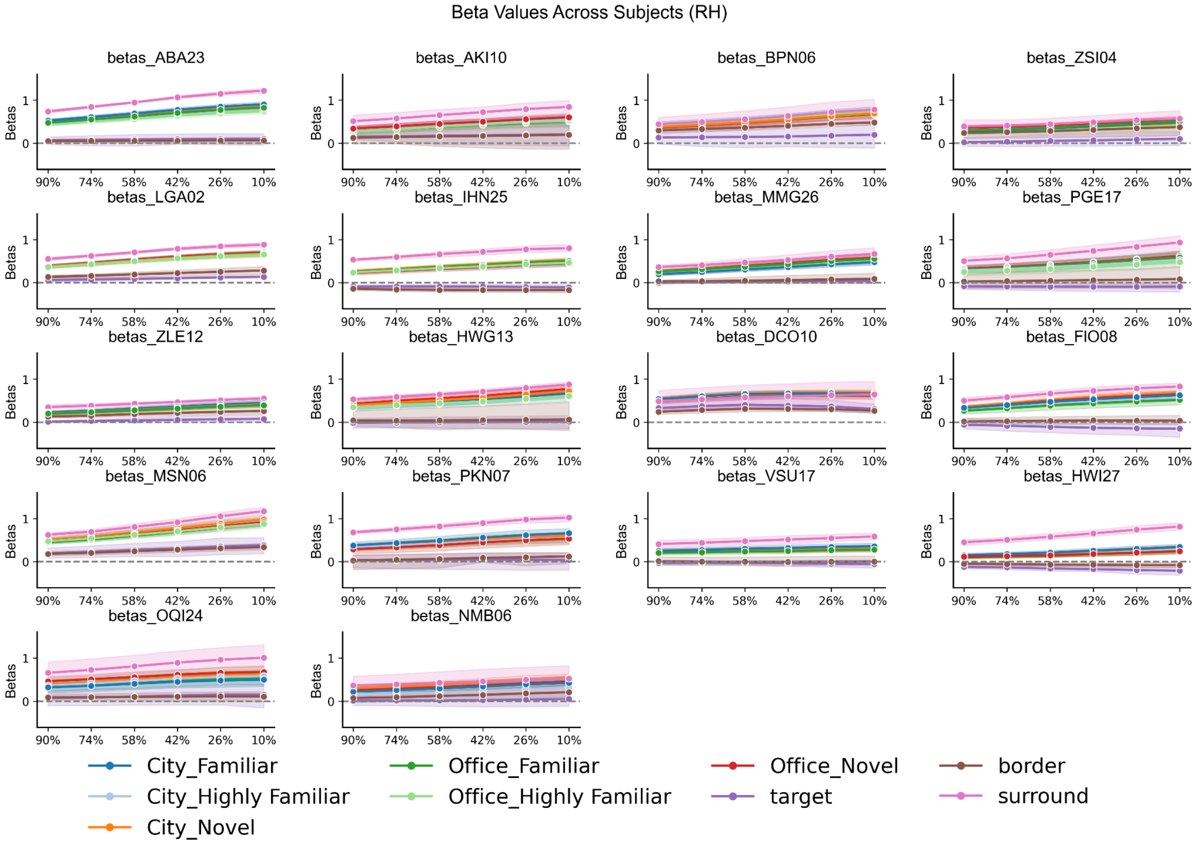


**Supplementary Fig. 3. Individual univariate BOLD betas across all layers in the V1 feedforward area in Experiment 1 (N = 18).** Shown are mean beta estimates for the three mapping conditions (surround, border and target) and the six experimental conditions: City Novel, City Familiar, City Highly Familiar, Office Novel, Office Familiar and Office Highly Familiar.

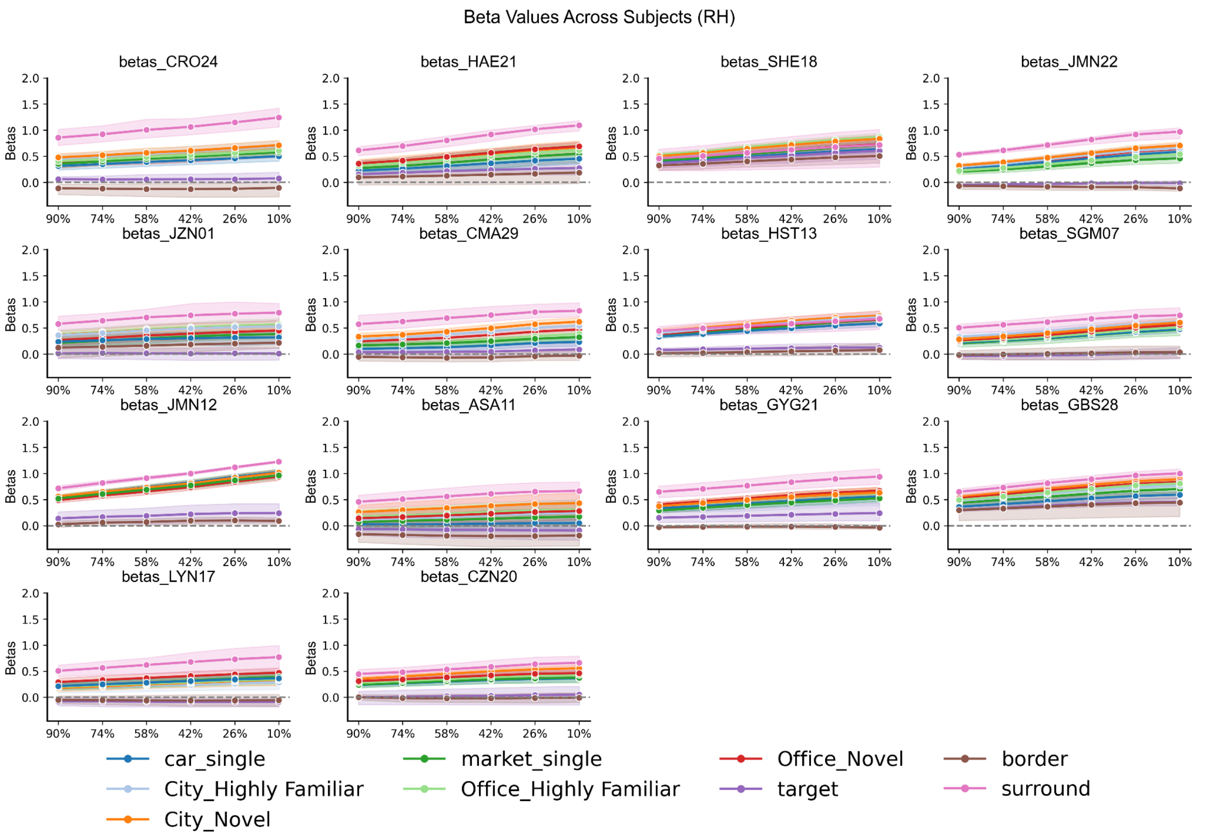


**Supplementary Fig. 4 Individual univariate BOLD betas across all layers in the V1 feedforward area in Experiment 2 (N = 14)**. Shown are mean beta estimates for the three mapping conditions (surround, border and target) and the six experimental conditions: City Novel, car-single (surround completion image 1), City Highly Familiar, Office Novel, market-single (surround completion image 2) and Office Highly Familiar.

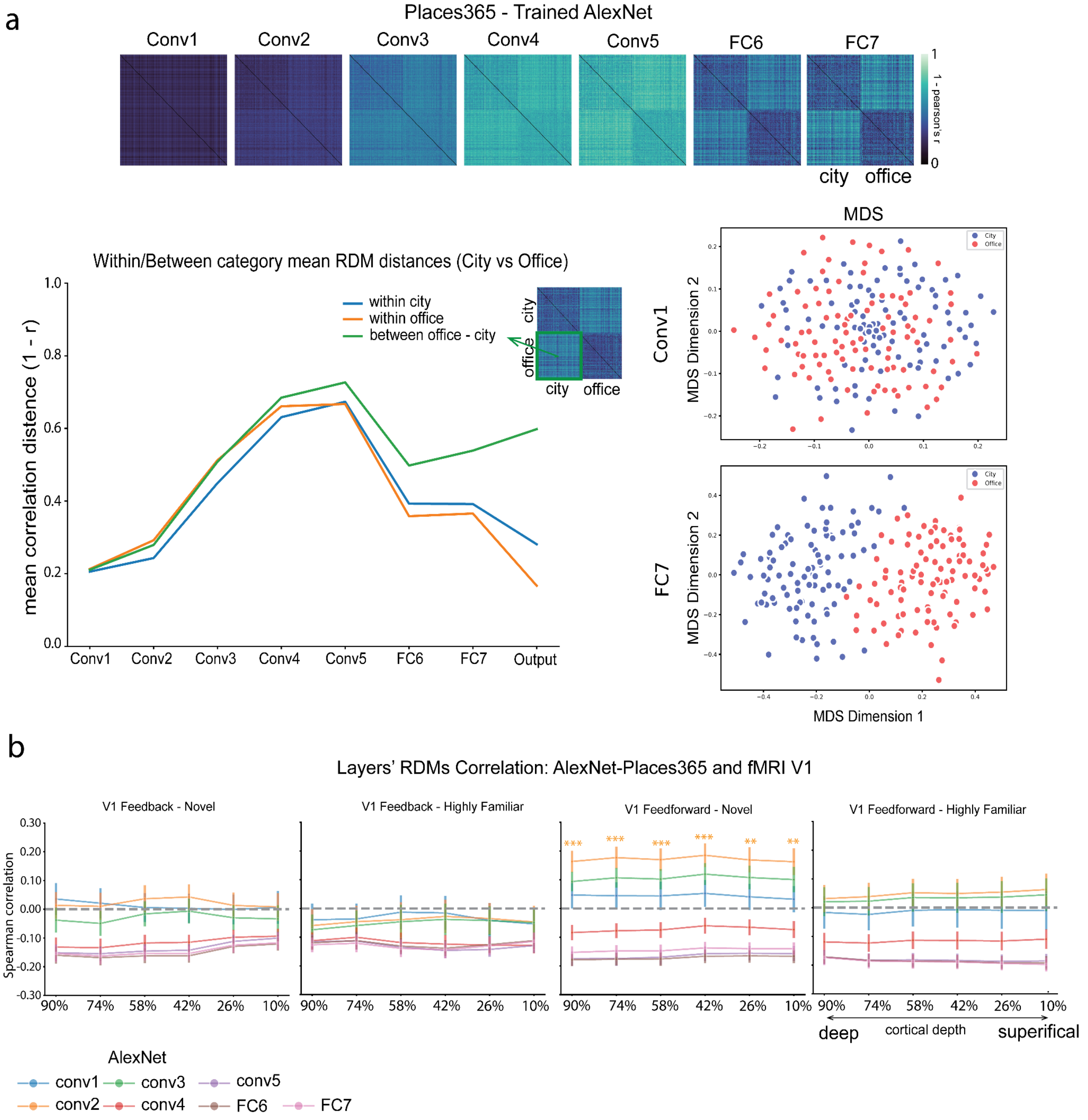


**Supplementary Fig. 5. AlexNet representational geometry and its correspondence with laminar V1 representations. a.** (1 - Pearson's r) representational dissimilarity matrices (RDMs) of AlexNet model features from Conv1 to FC7. The line plot shows the mean dissimilarity values within the city (blue), within the office (orange), and between the city and office (green) across each layer. The multidimensional scaling (MDS) plots illustrate the feature-based separation between city and office images in the conv1 (lower-level features) and FC7 (higher-level, more abstract) layers. **b.** Spearman correlations between AlexNet layer 8 x 8 category-run RDMs and fMRI layer 8 x 8 category-run RDMs in the Novel and Highly Familiar conditions, separately shown for V1 feedback and feedforward regions.

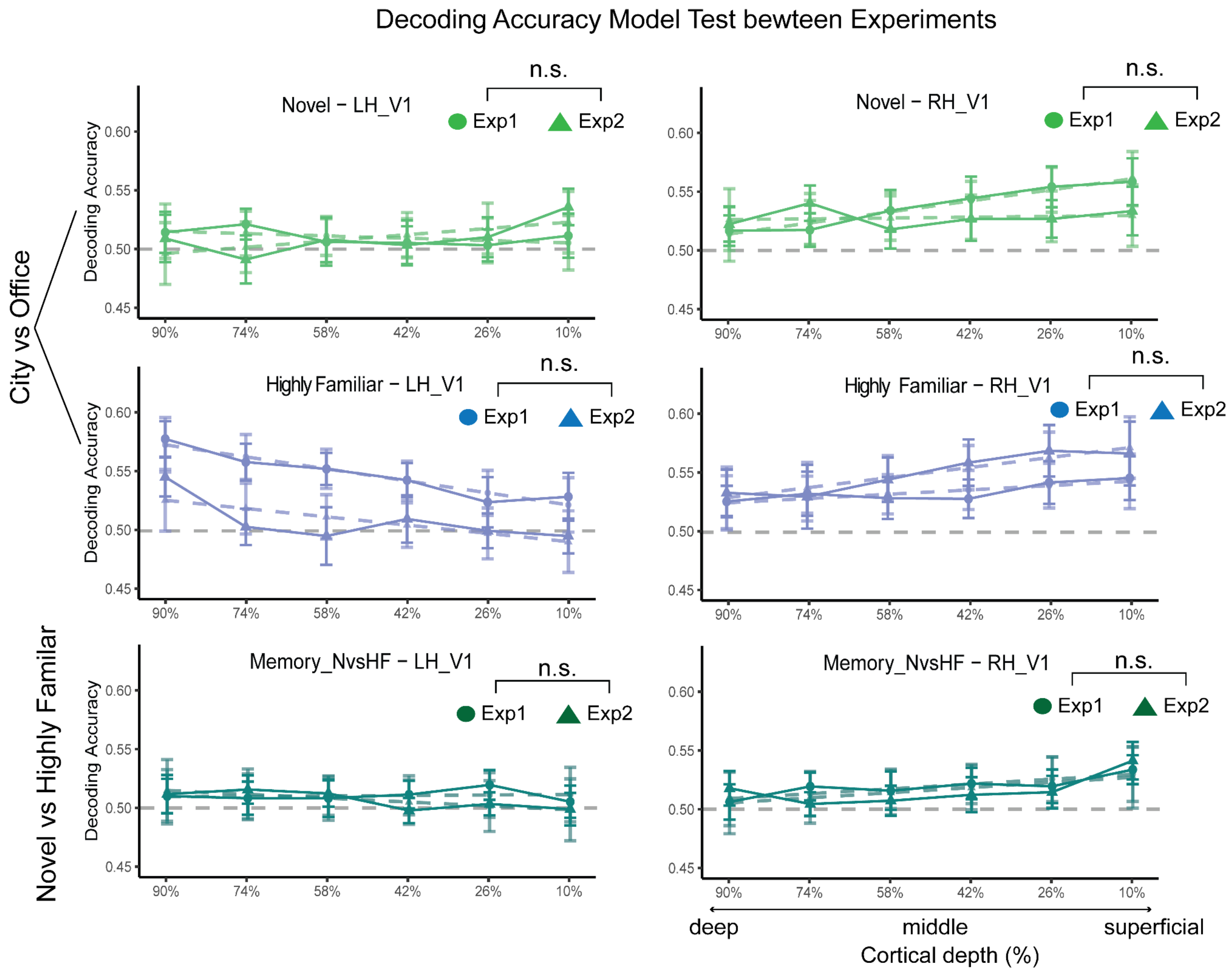


**Supplementary Fig. 6. Cross-experiment decoding and model-predicted accuracy profiles across cortical depth.** Decoding accuracy and model test across cortical depth in Experiments 1 and 2. Decoding accuracy is plotted across cortical depths from deep (90%) to superficial (10%). The blue solid line represents decoding accuracy for the Highly Familiar-categorical condition, the green solid line for the Novel-categorical condition, and the dark green solid line for the Novel versus Highly Familiar memory condition. The dashed lines indicate predicted values from the linear mixed-effects model. Circles dot means data from Experiment 1, and triangles denote data from Experiment 2. The chance level is 0.5 (50%).

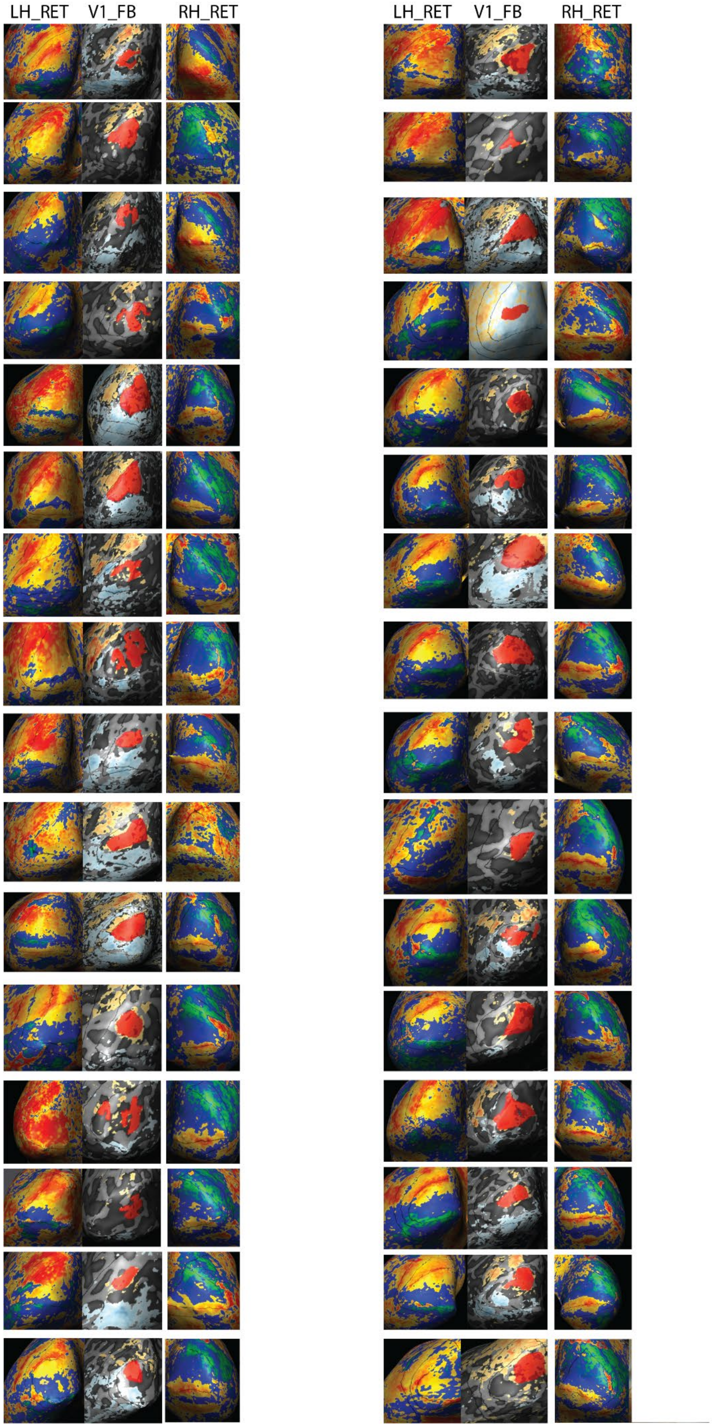


**Supplementary Fig. 7. Retinotopy and ROI definition for individual participants in both experiments (N = 32).** The V1 boundary was defined using retinotopic polar-angle mapping, and ROIs were identified from mapping trials for both V1 feedback and feedforward regions. The feedback ROI was located in the left hemisphere (LH), corresponding to the occluded lower-right visual quadrant (red).

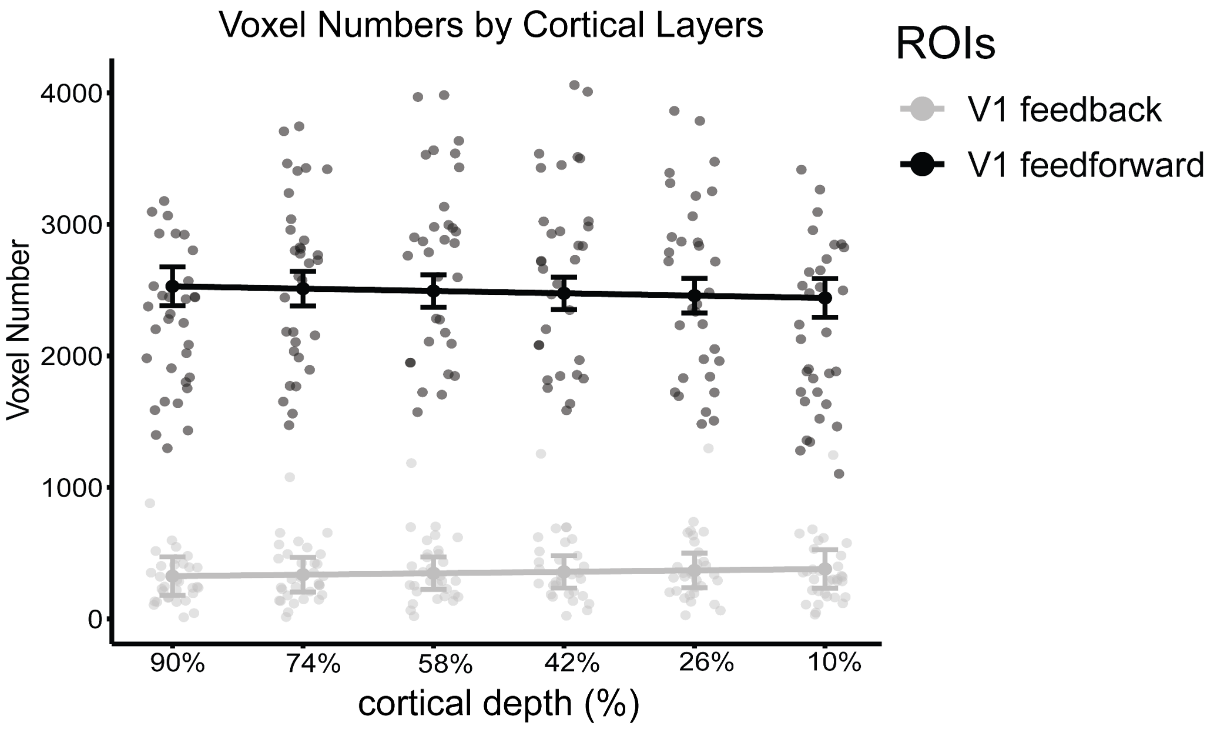


**Supplementary Fig. 8. Voxel counts across cortical depth in V1 feedforward and feedback ROIs.** Voxel number model test: the figure shows voxel numbers in V1 feedforward (black) and V1 feedback (grey), with lines (model predictions) and dots (actual data). None of the layer trends was significant (all $p > 0.05$), suggesting that voxel numbers did not differ across cortical depths in each ROI.

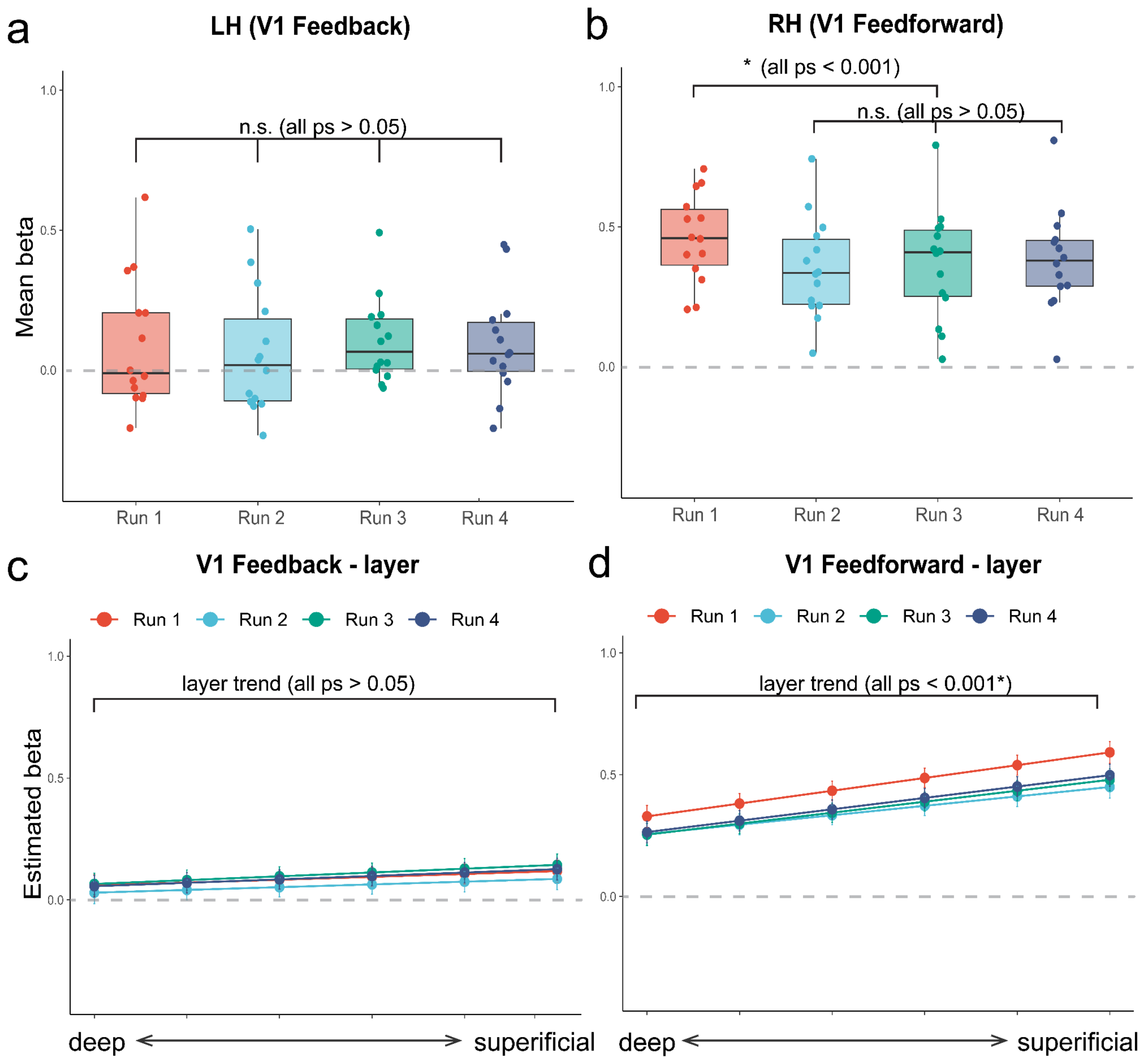


**Supplementary Fig. 9. Run-wise BOLD responses in the surround-completion condition (Exp2, N = 14). a, b.** Mean BOLD beta estimates across the four scanning runs in the V1 feedback ROI **(a)** and V1 feedforward ROI **(b)**. Points represent individual participants. **c, d.** Model-estimated BOLD responses across cortical depths (deep to superficial) for each run in the V1 feedback **(c)** and V1 feedforward **(d)** ROIs. Error bars indicate standard errors. *$p < 0.05$, Benjamini–Hochberg false discovery rate corrected.

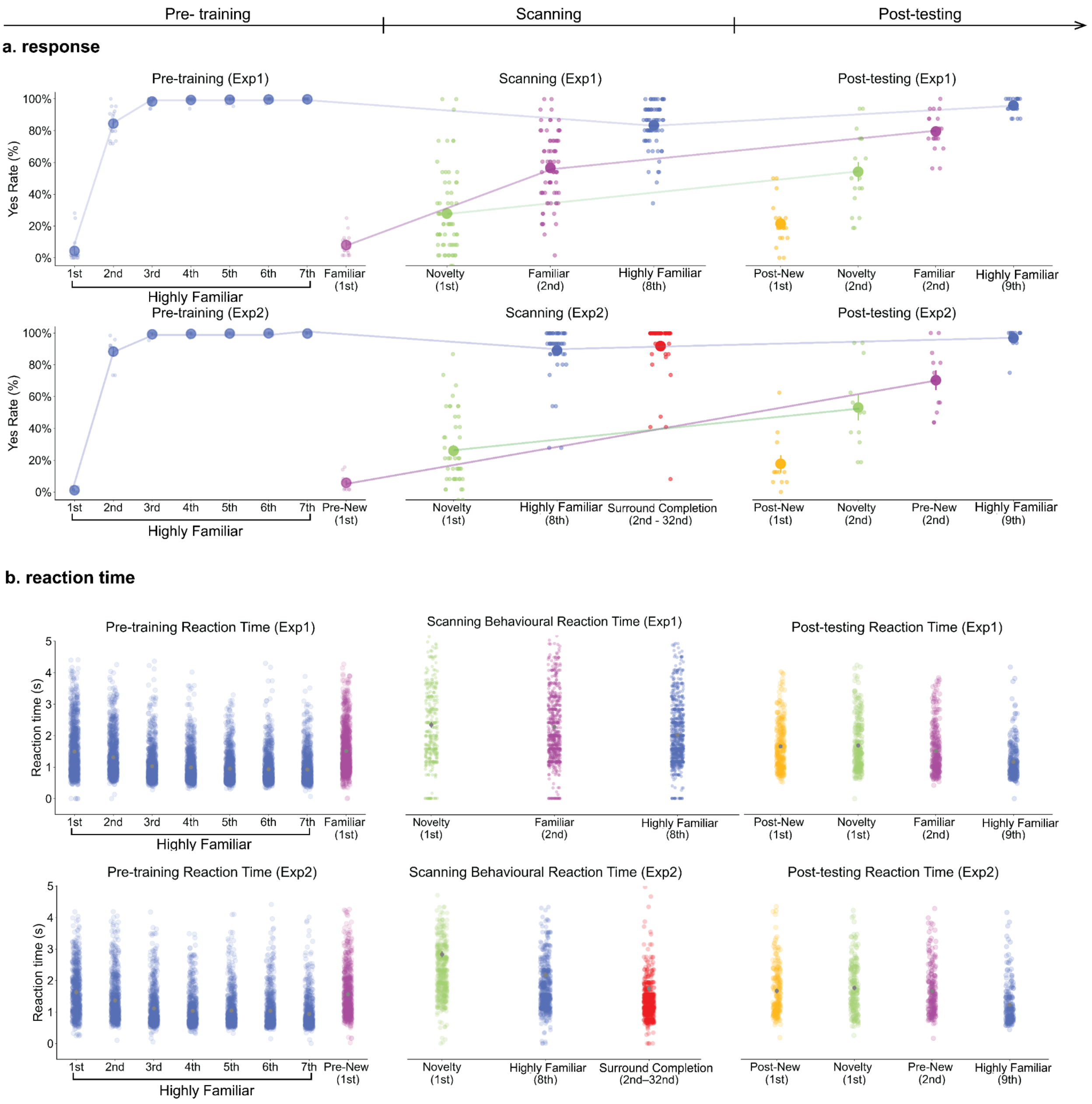


**Supplementary Fig. 10. Behavioural responses and reaction time across pre-training, scanning, and post-testing sessions in both Experiments. a.** Participants' response 'yes' rates and reaction times across pre-training, scanning, and post-testing sessions for each experiment. In the pre-training session, responses are shown for the Highly Familiar condition (both experiments; blue), the Familiar condition (Experiment 1; purple), and the Pre-New condition (Experiment 2; purple). The scanning session included Novel (green), Highly Familiar (blue), Familiar (Experiment 1; purple), and Single-Repeated (Experiment 2; red, data from 2nd to 32nd presentations). The post-testing session included Post-New (yellow), Novel (green), Familiar/Pre-New (purple), and Highly Familiar (blue). **b**. Reaction time across pre-training, scanning, and post-testing sessions in both Experiments

**Supplementary Note 1 (behavioural analysis):**

Experiment 1 included 18 participants in total. Behavioural response data were available from 18 participants in the pre-training session, 17 participants in the scanning session, and 18 participants in the post-test session. Experiment 2 included 14 participants in total. Behavioural response data were available from 13 participants in the pre-training session, 12 participants in the scanning session, and 14 participants in the post-test session. The reduced number of available behavioural datasets in some sessions was due to technical problems (MRI-compatible button box and stim-pc). Across all sessions, participants performed an old/new recognition task, in which they indicated whether each image had been seen previously at that point in the experiment. In Experiment 1, during the pre-training session, Highly Familiar images were presented seven times; the correct response was “no” on the first presentation and “yes” from the second to the seventh presentation. Familiar images were presented once during pre-training and were therefore new at that point, so the correct response was “no”. During the scanning session, Novel images had never been seen before, and the correct response was “no”, whereas Familiar and Highly Familiar images had been seen previously and the correct response was “yes”. During the post-test session, Novel, Familiar and Highly Familiar images had all been presented previously during the experiment, and the correct response was therefore “yes”, whereas post-new images were newly introduced at post-test and the correct response was “no”. Experiment 2 used the same response-coding logic for Novel and Highly Familiar images, but the Familiar condition was replaced by a pre-new condition during training. These pre-new images were presented once during the pre-training session and were new at that point, so the correct response was “no”; at post-test, they had been seen previously, and the correct response was therefore “yes”. In the scanning session of Experiment 2, the surround-completion condition consisted of repeated presentations of a single occluded image. The first presentation was new, and the correct response was “no”, whereas from the second to the thirty-second presentation the image had been seen previously and the correct response was “yes”. Reaction times were also calculated for trials with valid button responses in both experiments. Statistical analyses were conducted in Python v.3.11 using pandas, NumPy, SciPy, statsmodels, pingouin and scikit-posthocs. Response rates and reaction times were analysed after assessing distributional assumptions. Normality was evaluated using the Shapiro–Wilk test, and equality of variance was assessed using Levene's test. When both assumptions were satisfied, condition effects were tested with one-way ANOVA, followed by Tukey's honestly significant difference post hoc comparisons. If the normality assumption was satisfied but the assumption of equal variance was violated, Welch's ANOVA was used, followed by Games–Howell post hoc tests. When the normality assumption was violated, non-parametric Kruskal–Wallis tests were performed, followed by pairwise Dunn's tests with Benjamini–Hochberg correction for multiple comparisons. All statistical tests were two-tailed, and significance was defined as $P < 0.05$. These analyses were applied to behavioural data from all three sessions in both experiments. Data were visualised using matplotlib and seaborn.

**Experiment 1:**

**Recognition task (response):**

**Pre-training:** Shapiro-Wilk tests indicated that the assumption of normality was violated for most sessions ($ps < 0.05$, except for the 1st presentation), and Levene's test showed significant heterogeneity of variances across conditions ($p < 0.001$). Therefore, a Kruskal-Wallis test was performed to examine differences in the rate of "Yes" button presses across sessions. The test revealed a significant main effect of condition, $H(7) = 117.13$, $p < 0.001$. Dunn's post-hoc tests (BH-corrected) showed that "Yes" response rates increased significantly from the early (1st-2nd) to later (3rd-7th) sessions (all $p < 0.001$). The Familiar condition did not differ from 1st or 2nd ($p > 0.05$), but it did differ significantly from all later sessions ($p < 0.001$), indicating a strong session-dependent modulation in affirmative response rates.

**Scanning:** Shapiro–Wilk tests indicated that the normality assumption was met for all conditions: Familiar, Novel, and Highly Familiar. However, Levene's test indicated unequal variances across groups ($p = 0.0405$). Therefore, a Welch's ANOVA was conducted to compare the rate of pressing the "Yes" button among the Novel, Familiar, and Highly Familiar conditions. The analysis revealed a significant main effect of condition, $F(2, 28.31) = 62.37$, $p < 0.001$. Games–Howell post hoc comparisons showed that all conditions differed significantly from one another. The "Yes" response rate was significantly higher for Familiar than Novel trials, $p < 0.001$, and significantly higher for Highly Familiar than both Familiar and Novel trials, both $p < 0.001$.

**Post-testing:** Shapiro-Wilk tests indicated that the data for the Post-New condition deviated from normality ($p = 0.0002$), while other conditions did not (all ps > 0.1), and Levene's test revealed unequal variances across conditions ($p = 0.0001$). Consequently, a Kruskal-Wallis test was conducted to compare the rate of "Yes" button presses across the four conditions. The test revealed a significant main effect of the condition on "Yes" response rate, $H(3) = 54.05$, $p < 0.001$. Dunn's post-hoc tests (BH-corrected) showed that the "Yes" response rate differed significantly between most conditions. Specifically, Post-New and Highly Familiar conditions differed considerably from all other conditions (all $p < 0.001$). A smaller but significant difference was also observed between Familiar and Novel ($p = 0.036$).

**Reaction time:**

**Pre-training:** Shapiro-Wilk tests indicated significant deviations from normality for all conditions (all ps < 0.001), and Levene's test showed unequal variances ($p < 0.001$). Therefore, a Kruskal–Wallis test was conducted to examine differences in reaction time across sessions. The test revealed a significant main effect of condition, $H(7) = 1429.82$, $p < 0.001$. Dunn's post-hoc tests (BH-corrected) showed that reaction times differed significantly across most session pairs. Reaction times decreased substantially from the 1st to later presentations of the Highly Familiar condition (all $p < 0.001$), and the Familiar condition also differed significantly from all sessions (all $p < 0.01$). No significant differences were found among the final sessions (5th-7th, all $p > 0.08$), suggesting that reaction times plateaued with continued exposure.

**Scanning:** Shapiro-Wilk tests indicated that the reaction time data deviated from normality in all conditions (all $p < 0.001$), and Levene's test showed unequal variances across conditions ($p < 0.001$). Therefore, a Kruskal-Wallis test was used to compare reaction times among the three familiarity conditions. The test revealed a significant main effect of familiarity on reaction time, $H(2) = 27.0496$, $p < 0.001$. Dunn's post-hoc tests (BH-corrected) showed that reaction times for Highly Familiar stimuli were significantly higher than those for Familiar and Novel conditions (both ps < 0.001). No significant differences were found between Novel and Familiar conditions (p > 0.8).

**Post-testing:** Shapiro-Wilk tests indicated that reaction times in all conditions deviated from normality (all ps < 0.001), and Levene's test showed unequal variances across conditions (p < 0.001). Therefore, a Kruskal-Wallis test was conducted to assess differences in reaction times across the four conditions. The test revealed a significant main effect of condition, H(3) = 110.90, p < 0.001. Dunn's post-hoc tests (BH-corrected) showed that reaction times in the Highly Familiar condition differed significantly from all other conditions (all p < 0.001). Reaction times in the Pre-New condition were also significantly different from both Novel (p = 0.006) and Post-New (p = 0.007). No significant difference was observed between Novel and Post-New (p = 0.90).

**Experiment 2:**

**Recognition task (response):**

**Pre-training**: Shapiro–Wilk tests indicated that the normality assumption was violated for all conditions, including Pre-New and the first to seventh presentations of the Highly Familiar images, all ps < 0.05. Levene's test also indicated significant heterogeneity of variance across conditions (p = 0.0001). Therefore, a non-parametric Kruskal–Wallis test was used to compare the rate of pressing the "Yes" button across the eight conditions. The analysis revealed a significant main effect of condition, H(7) = 68.06, p < 0.001. Dunn's post-hoc comparisons with FDR (BH) correction showed that "Yes" response rates increased across repeated presentations. Specifically, Pre-New and the first presentation did not differ significantly from each other, p = 0.683, and neither differed significantly from the second presentation, both ps > 0.05. However, Pre-New and the first presentation differed significantly from the third to seventh presentations (all ps < 0.001). The second presentation also differed significantly from the third to seventh presentations (all ps < 0.05). No significant differences were found among the later presentations from the third to seventh presentation (all ps > 0.05).

**Scanning**: Shapiro–Wilk tests indicated that the normality assumption was violated for the Highly Familiar and Single conditions (both ps < 0.001), whereas the Novel condition did not significantly deviate from normality (p = 0.141). Levene's test indicated that the homogeneity of variance assumption was met ($p = 0.269$). Because the normality assumption was violated, a non-parametric Kruskal–Wallis test was used to compare “Yes” response rates across the Novel, Highly Familiar and Single conditions. The analysis revealed a significant main effect of condition, $H(2) = 23.46$, $p < 0.001$. Dunn’s post-hoc comparisons with Benjamini–Hochberg correction showed that the Novel condition differed significantly from both the Highly Familiar condition ($p = 0.001$) and the Surround completion condition ($p < 0.001$). No significant difference was observed between the Highly Familiar and Surround completion conditions ($p = 0.197$).

**Post-testing**: Shapiro-Wilk tests indicated that the data from the Pre-New ($p = 0.0047$) and Post-New ($p < 0.001$) conditions significantly deviated from normality, and Levene’s test showed unequal variances across conditions ($p = 0.0064$). Therefore, a Kruskal-Wallis test was conducted. The test revealed a significant effect of condition on ‘yes’ rate, $H(3) = 33.97$, $p < 0.001$. Dunn’s post-hoc tests (BH-corrected) showed that the rate of pressing the ‘yes’ button in the Post-New condition was significantly different from Novel ($p = 0.031$), Pre-New ($p = 0.001$), and Highly Familiar ($p < 0.001$). Accuracy in the Highly Familiar condition was also significantly higher than in the Novel ($p = 0.001$) and Pre-New ($p = 0.032$) conditions. No significant difference was found between Novel and Pre-New ($p = 0.23$).

**Reaction time:**

**Pre-training:** Shapiro–Wilk tests indicated that reaction times across all eight conditions, including Pre-New and the first through seventh presentations of the Highly Familiar images, significantly deviated from normality (all ps $< 0.001$). Levene’s test also indicated significant heterogeneity of variance across conditions ($p < 0.001$). Therefore, a non-parametric Kruskal–Wallis test was used to compare reaction times across conditions. The analysis revealed a significant main effect of condition on reaction time, $H(7) = 1020.11$, $p < 0.001$. Dunn’s post hoc comparisons with FDR (BH) correction showed that reaction times differed significantly across repeated presentations. Pre-New and the first presentation did not differ significantly ($p = 0.324$), but both differed significantly from the second through seventh presentations (all ps $< 0.001$). The second presentation also differed significantly from later presentations, all ps $< 0.001$. In addition, reaction times were not fully stable across later repetitions: the seventh presentation differed significantly from the third to sixth presentations, all ps $< 0.001$. These results indicate that reaction times changed systematically across repeated exposure, with a marked difference between the initial/new presentations and subsequent repeated presentations.

**Scanning:** Shapiro–Wilk tests revealed significant deviations from normality for all three conditions (all ps $< 0.001$), and Levene's test indicated unequal variances across groups ($p < 0.001$). Therefore, a non-parametric Kruskal–Wallis test was used to compare reaction times across the Novel, Highly Familiar and Surround completion conditions. The analysis revealed a significant main effect of condition on reaction time, $H(2) = 389.00$ ($p < 0.001$). Dunn's post-hoc comparisons with Benjamini–Hochberg correction indicated that reaction times differed significantly between all pairs of conditions (Novel > Highly Familiar > Surround completion, all ps $< 0.001$).

**Post-testing**: Shapiro-Wilk tests indicated significant departures from normality in all conditions (all ps $< 0.001$), and Levene's test indicated unequal variances across conditions ($p = 0.0002$). Therefore, a Kruskal-Wallis test was used. The analysis revealed a significant effect of condition on reaction time, $H(3) = 86.24$, $p < 0.001$. Dunn's post hoc tests (BH-corrected) showed that reaction time in the Highly Familiar condition was significantly different from Novel ($p < 0.001$), Pre-New ($p < 0.001$), and Post-New ($p < 0.001$) conditions. However, no significant differences were found among Novel, Pre-New, and Post-New (all ps $> 0.05$).
